# Human receptive endometrial assembloid for deciphering the implantation window

**DOI:** 10.1101/2023.07.27.550771

**Authors:** Yu Zhang, Rusong Zhao, Chaoyan Yang, Jinzhu Song, Peishu Liu, Yan Li, Boyang Liu, Tao Li, Changjian Yin, Minghui Lu, Zhenzhen Hou, Chuanxin Zhang, Zi-Jiang Chen, Keliang Wu, Han Zhao

**Author notes:** These authors contributed equally: Yu Zhang, Rusong Zhao. **Classifications:** Cell Biology.

## Abstract

Human endometrial receptivity is a critical determinant of pregnancy success; however, in vivo studies of its features and regulation are particularly challenging due to ethical restriction. Recently, the development of human endometrial assembloids has provided a powerful model to investigate this intricate biological process. In this study, we established a specialized human window-of-implantation (WOI) endometrial assembloid system that mimics the in vivo receptive endometrium. It not only reproduces the structural attributes of pinopodes and cilia, but also molecular characteristics of mid-secretory endometrium. Furthermore, the WOI endometrial assembloid exhibits hormone responsiveness, an energy metabolism profile characterized by larger and functionally enhanced mitochondria, increased ciliary assembly and motility, and epithelial-mesenchymal transition (EMT), as well as promising potential for embryo implantation. As such, WOI assembloids hold great promise as a platform to unravel the intricate mechanisms governing the regulation of endometrial receptivity, maternal-fetal interactions, and associated pathologies, ultimately driving impactful advancements in the field.

## Introduction

The human endometrium, a complex tissue comprising diverse cell types, undergoes shedding (menstrual phase), regeneration (proliferative phase), and differentiation (secretory phase) under the coordinated regulation of estrogen and progesterone throughout the menstrual cycle (1). A brief interval, termed the “window-of-implantation (WOI)” or mid-secretory phase, peimits embryo implantation into the endometrium (1). The endometrium at the site of embryo implantation harbors a heterogeneous population of cells, including ciliated epithelial cells, decidualized stromal cells and immune cells. Cyclic endometrial changes analogous to those observed in humans are exclusive to apes, Old World monkeys, molossid bats, and spiny mice, but are absent in conventional laboratory mice (1). This renders typical mouse models inadequate for accurately recapitulating human endometrium. Existing endometrial cell lines, such as Ishikawa and HEEC cells, are composed of a single cell type, and thus fail to reproduce the intricate physiological architecture and function of the endometrium.

Replicating and reconstructing human organs has become indispensable for elucidating tissue physiology and function. Organoid, a self-assembled 3D structure, closely resemble in vivo tissue or organ (2). They offer high expansibility, alongside conserved phenotypic, and functional properties, emerging as powerful tools for investigating tissue physiology and disease (3). In 2017, the first long-term and hormone-responsive human endometrial organoid was derived from adult stem cells in endometrial biopsies (2, 3). Based on this, Margherita Y. Turco, et al generated endometrial organoids from menstrual effluent using non-invasive methods (4), while Takahiro Arima et al. engineered polarity-reversed organoids to study embryo implantation(5). Beyond adult stem cells, pluripotent stem cells were also induced to endometrial stromal fibroblasts and epithelium, and then cocultured to form organoids(6, 7), which exhibited vigorous proliferative capacity but lacked immune cells and other components of the microenvironment. Several studies have additionally incorporated immune cells into endometrial organoid co-culture systems(8). To recapitulate the full spectrum of human endometrial cell types in vitro, endometrial assembloids have evolved from epithelial organoids(9), to assemblies of epithelial and stromal cells(10, 11) and then to stem cell-laden 3D artificial endometrium(12, 13), which were solely closer but not completely identical to the endometrium. Additionally, pathological endometrial organoid models have been established for conditions like endometriosis, endometrial hyperplasia, Lynch syndrome, and endometrial cancer (14). These organoids faithfully simulated endometrial morphology, hormone responsiveness, and physiological and pathological processes in vitro (3, 9, 15–17), facilitating the study of physiological phenomena (15, 18, 19), pathogenic mechanisms(16) and drug screening(14).

Although various regulators are implicated in the WOI, our understanding of embryo implantation during this period remains limited by ethical constraints and a paucity of in vitro receptive endometrial models. The transition from proliferative to receptive endometrium involves dynamic processes, including decidualization, epithelial-mesenchymal transition (EMT) and ciliated epithelial development (1, 20), which most in vitro models have yet to fully recapitulate. Here, we established a hormone-regulated receptive endometrial assembloid system that recapitulates in vivo WOI features, providing a platform to study physiological/pathological endometrial function and maternal–fetal interactions.

## Results

### Developing receptive endometrial assembloids in vitro

To generate endometrial assembloids in vitro, pre-receptive endometrium from reproductive-age women was dissociated into single cells or small cell masses. These cells then self-assembled into assembloids induced by various small molecules, such as Noggin, EGF, FGF2, WNT-3A and R-Spondin1 in expansion medium (ExM) (Fig.1A∼B, Fig.S1A). The assembloids derived from the first three generation are used for experiments (Fig.S1B). The endometrial assembloids comprised vesicle-like glands and fibrous stromal cells (Fig. S1C, Video S1∼S4). The epithelium marker E-cadherin, stromal cell marker vimentin, and endometrial gland marker FOXA2 were confirmed in cultured assembloid, which exhibited morphological similarity to the endometrium (Fig. S1D). Moreover, the endometrial assembloids exhibited substantial expression of the proliferation marker Ki67, with no detectable cleaved caspase-3 (apoptosis marker), indicative of strong proliferative capacity (Fig. S1E).

**Fig 1.**
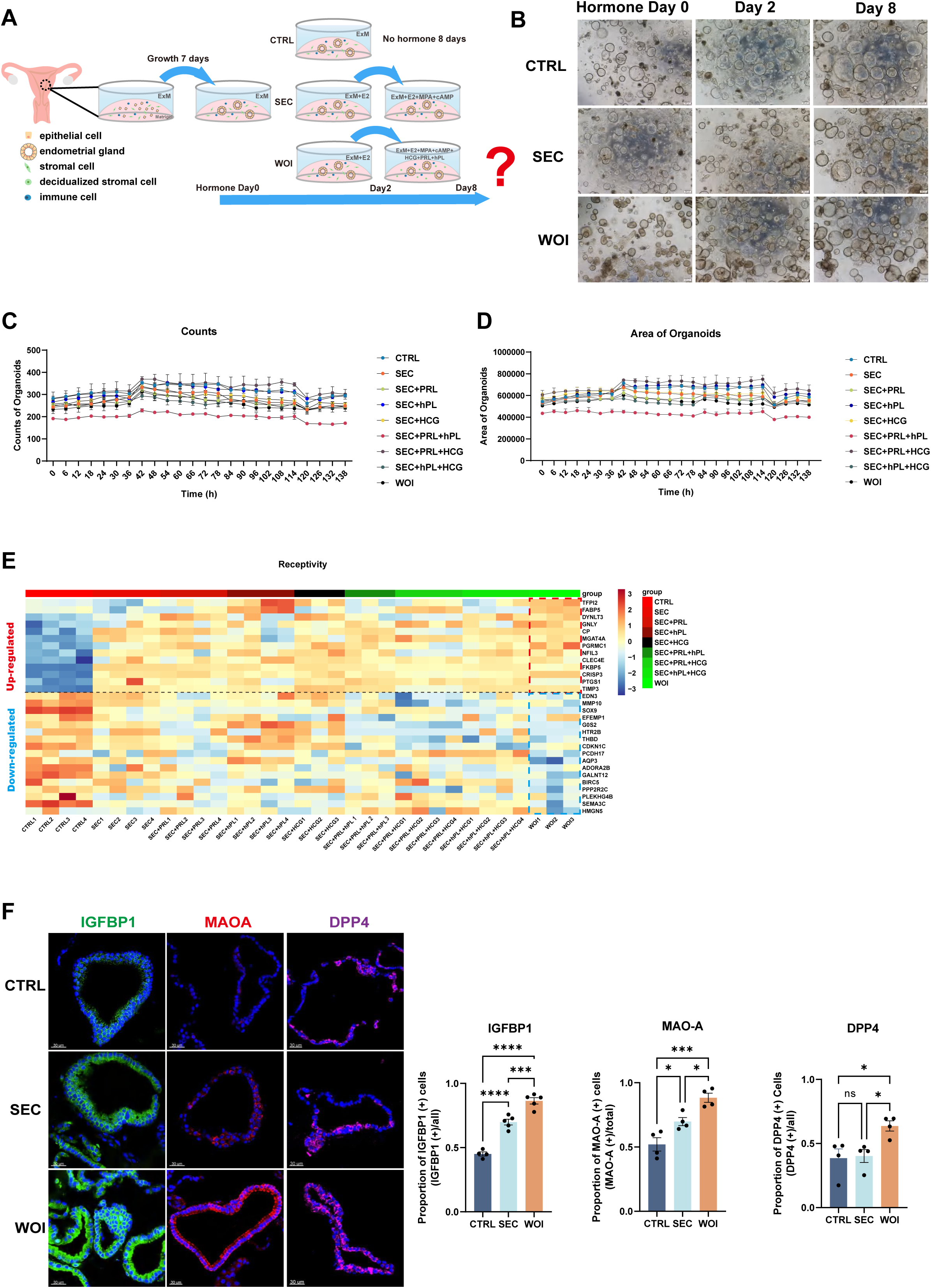
Developing receptive endometrial assembloids in vitro (A) Human endometrial assembloids constructed from adult stem cells were treated with expansion medium (ExM) (CTRL) or subjected to hormonal stimulation. Timeline of endometrial assembloid cultured by ExM (CTRL), ovarian steroid hormones simulating secretory phase (SEC), ovarian steroid hormones combining PRL and placental hormones to mimic the window of implantation (WOI). (B) Endometrial assembloids from the CTRL, SEC, and WOI groups, which were subjected to hormone treatment on Days 0, 2, and 8, exhibited comparable growth patterns throughout the culture period. Scale bar = 200 μm. (C) The dynamic changes of the counts of assembloids over time in each hormone regimen. (D) The dynamic changes of the area of assembloids over time in each hormone regimen. (E) Heatmap showing receptivity related gene expression profile of assembloids in each hormone regimen. The color represents log-transformed fold change of gene expression. (F) Validation of receptivity markers (IGFBP1, MAOA and DPP4) with immunofluorescence (IF) in the CTRL, SEC and WOI endometrial assembloids in vitro. Nuclei were counterstained with DAPI. Scale bar = 30 μm. The bar chart displaying the quantitative comparison of receptivity markers among three groups. *P ≤ 0.05, **P ≤ 0.005, ***P ≤ 0.0005, ****P ≤ 0.0001.

Functionally, endometrial assembloids effectively secreted glycogen into the lumen, mirroring the nutrient-secreting activity of the endometrium to support embryo implantation (Fig. S1D). Moreover, after two days of estrogen (E2) treatment followed by fourteen days of medroxyprogesterone acetate (MPA) and cAMP, the assembloids exhibited significant transcriptional upregulation of progesterone receptor (PRA/B), a modest increase in estrogen receptor α (ERα), and enhanced expression of estrogen-responsive genes *EGR1* and *OLFM4*, along with progesterone-responsive genes *PGR* and *PAEP* (Fig. S1F∼S1G). It reflected hormonal responsiveness analogous to that of the in vivo endometrium (2).

To identify hormone regimens that induce implantation window, pregnancy-related hormones were supplemented into the culture system following 2 days of E2 priming. E2 and MPA drive the transition of endometrial assembloids to the secretory phase, while pregnancy hormones promote further differentiation. Prolactin (PRL) promotes immune regulation and angiogenesis during implantation(2, 21). Human chorionic gonadotropin (hCG) improves endometrial thickness and receptivity (22, 23). Human placental lactogen (hPL) promotes the development and function of endometrial glands(24). Hormone dosages were primarily based on peri-pregnant maternal systemic or local endometrium levels (2). Multigroup comparison revealed similar counts, area, and average intensity of assembloids over time (Fig.1C-D, Fig.S1H). However, only the final cocktail (i.e., a combination of E2, MPA, cAMP, PRL, hCG and hPL) exhibited an endometrial receptivity-related gene expression profile, with high expression of receptivity-promoting genes and low expression of receptivity-inhibiting genes relative to other hormone formulations (Fig.1E). The assembloids generated with this regimen were defined as WOI assembloids (Fig. 1A). For controls, assembloids maintained in ExM served as the “control (CTRL)” group, while those treated with 2 days of E₂ followed by 6 days of E₂, MPA, and cAMP were induced to the secretory phase (as previously described (9)) and designated the “secretory (SEC)” group (Fig. 1A). There was no significant difference in the morphology of assembloids among the three groups (Fig.1B), but WOI assembloids exhibited elevated expression of the receptivity markers IGFBP1, MAOA, and DPP4 (Fig.1F), and increased glycogen secretion (Fig.S1I). Theoretically, the WOI assembloids originate from the secretory phase and thus share characteristics with SEC assembloids, but crucially, they are expected to exhibit hallmark features of the mid-secretory phase.

### Receptive endometrial assembloids mimicked the implantation - window endometrium

Single-cell transcriptomics analysis, with reference to CellMarker, PanglaoDB, Human Cell Atlas, Human Cell Landscape, and scRNASeqDB, and prior endometrium related studies(1, 9, 11, 25), identified the presence of epithelial, stromal, and immune cells within WOI assembloids (Fig. 2A, Fig. S2A∼S2B). Comparative analysis of scRNA-seq data from our assembloids and mid-secretory endometrial tissue (as described by Stephen R. Quake et al. in 2020 (1) and Garcia-Alonso in 2021 (25)) revealed that WOI assembloids exhibited similarities to the mid-secretory endometrium in terms of glandular epithelium, ciliated epithelium, and stromal cells (Fig.2A, Fig.S2C∼S2F). The morphology of immune and stromal cells was visualized via 3D clearing staining and light sheet microscopy imaging, with vimentin labeling stromal cells (Vimentin^+^ or Vimentin^+^ F-actin^+^), CD45 and CD44 indicating immune cells, and FOXA2 identifying glands (Fig.2B∼2D). Furthermore, flow cytometry was employed to validate the presence and subset composition of immune cells. White blood cells (WBC) were identified as CD45^+^ cells, with T cells, macrophages and NK cells characterized as CD45^+^CD3^+^, CD45^+^CD68^+^CD11b^+^ and CD56^+^CD16^-^ cells, respectively (Fig. 2E). WOI assembloids displayed characteristic features of the receptive endometrium. On the one hand, they secreted significantly higher levels of glycogen into the lumen compared to other groups (Fig. S1I). On the other hand, they possessed various characteristic microstructure of implantation window, including elongated microvilli and increased glycogen, pinopodes, and especially cilia (Fig.2F).

**Fig 2.**
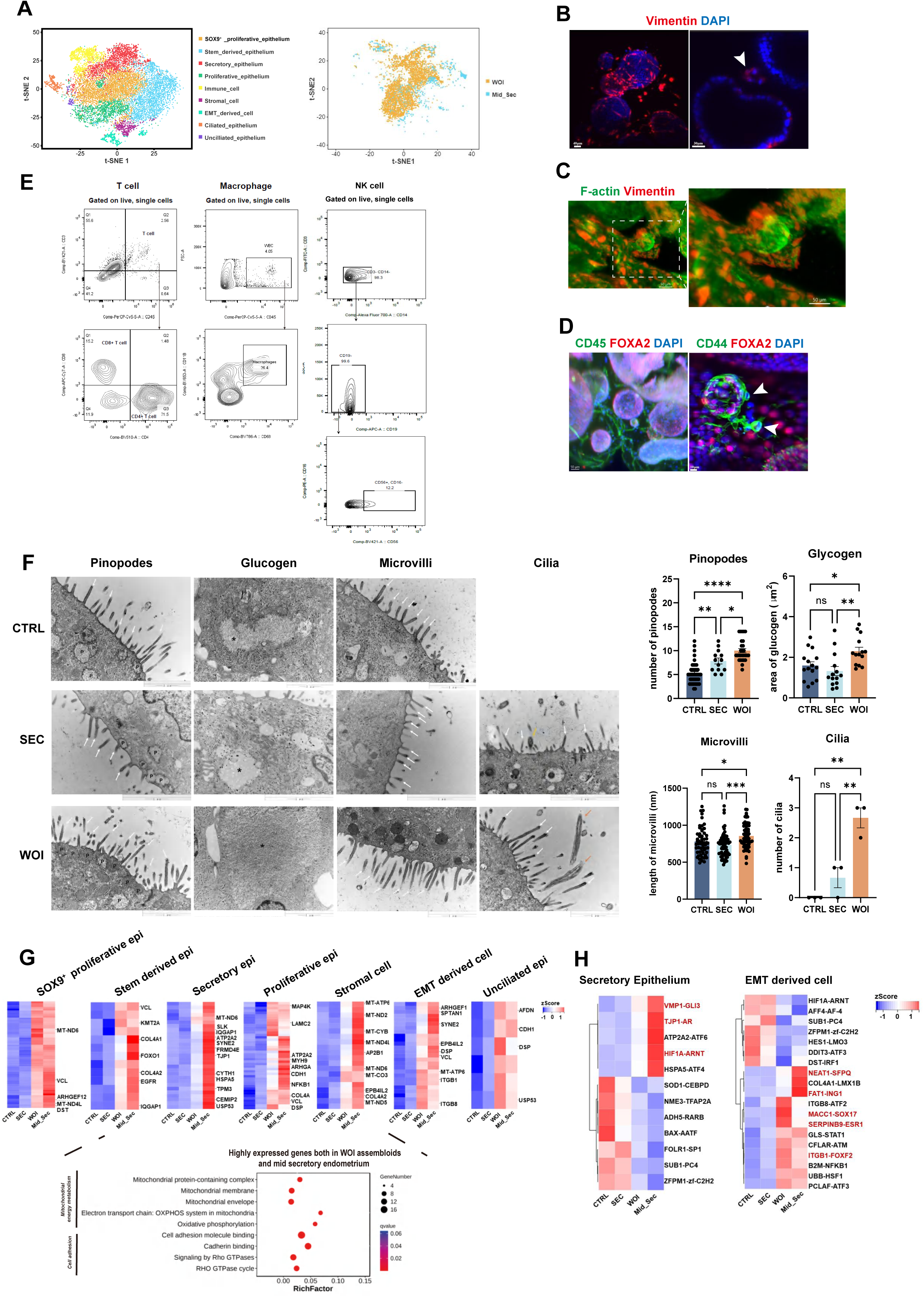
Receptive endometrial assembloids mimicked the implantation - window endometrium (A) T-SNE plot of scRNA-seq data from three individual endometrial assembloids of the CTRL, SEC and WOI groups (left). T-SNE plot of combined scRNA-seq data from the three kinds of assembloids and mid-secretory endometrium (right). (B) Exhibition of stromal cell marked by vimentin of CTRL assembloid through whole-mount clearing, immunostaining and light sheet microscopy imaging. Nuclei were counterstained with DAPI. The arrowhead indicates stromal cells. Scale bar = 40 μm (left), Scale bar = 30 μm (right). (C) Whole-mount immunofluorescence showed that Vimentin^+^ F-actin^+^ cells (stromal cells) were arranged around the glandular spheres that were only F-actin^+^. Scale bar = 50 μm. (D) Exhibition of immune cell marked by CD45 and CD44, and endometrial gland marked by FOXA2 of CTRL assembloid through whole-mount clearing, immunostaining and light sheet microscopy imaging. Nuclei were counterstained with DAPI. The arrowhead indicates immune cells. Scale bar = 50 μm (left), Scale bar = 10 μm (right). (E) Flow cytometric analysis of T cells and macrophages in the CTRL endometrial assembloid. Gating strategy used for determining white blood cells (WBC) (CD45^+^ cells), T cells (CD45^+^CD3^+^ cells) and macrophages (CD45^+^CD68^+^CD11b^+^ cells). (F) Electron micrograph of the CTRL (top), SEC (middle) and WOI (bottom) endometrial assembloid showing pinopodes (P), glycogen granule (asterisk), microvilli (white arrows) and cilia (orange arrows). Scale bar = 1 μm. Quantitative comparison of pinopodes, glycogen, microvilli, and cilia in the CTRL, SEC and WOI assembloids. *P ≤ 0.05, **P ≤ 0.005, ***P ≤ 0.0005, ****P ≤ 0.0001. (G) Heatmap and bubble diagram illustrating highly expressed genes as well as GO functions enriched in both assembloids during the WOI and mid secretory endometrial tissue in terms of SOX9^+^ proliferative epithelium, stem-derived epithelium, secretory epithelium, proliferative epithelium, unciliated epithelium, stromal cells and EMT-derived cells. The color of heatmap represents log-transformed fold change of gene expression. (H) Heatmaps showing differentially expressed TFs of endometrial assembloids and endometrium in the secretory epithelium (left) and EMT-derived cells (right). The color represents log-transformed fold change of gene expression.

We next focused on transcriptional profiling and regulation during the mid-secretory phase, as this phase marks the implantation window and involves substantial transcriptional alterations (Fig.2G∼2H). For this analysis, we referred scRNA-seq data of the mid-secretory phase from Stephen R. Quake 2020 (1). Pathways related to mitochondrial energy metabolism and cell adhesion were significantly upregulated in both WOI assembloid and mid-secretory endometrium, relative to CTRL and SEC assembloids (Fig. 2G). Furthermore, key transcription factors (TFs) associated with implantation—identified in secretory epithelial cells and EMT-derived cells—exhibited highly conserved expression patterns between WOI assembloids and the mid-secretory endometrium. Specifically, secretory epithelium shared comparable TFs linked to hypoxia response (e.g., HIF1A(26)), embryo implantation (e.g., FBLN1(27)), lipid metabolism (e.g., VMP1(28)), cell migration, and cell junction (e.g., TJP1(29)) (Fig. 2H). Similarly, EMT-derived cells also expressed conserved TFs involved in endometrial decidualization (e.g., S100A10(30)), EMT (e.g., FAT1(31) and FOXF2(32)), and receptivity (e.g., NEAT1(33), SERPINB9(34), SOX17(35) and SOX4(36)) (Fig. 2H).

To further validate the receptive phenotype of WOI assembloids, we performed Endometrial Receptivity Analysis (ERA), a gene expression profiling-based diagnostic assay that integrates high-throughput sequencing and machine learning to quantify the expression of endometrial receptivity-associated genes (37). Currently employed in clinical practice to assess endometrial receptivity and guide personalized embryo transfer, ERA revealed that WOI assembloids derived from pre-receptive endometrial tissue successfully transitioned to a receptive state (Fig. S1J).

Collectively, these findings confirm that WOI assembloids closely recapitulate the structural and molecular characteristics of the in vivo endometrium during the implantation window.

### Receptive endometrial assembloids recapitulate WOI-associated hormone response

To characterize the biological hallmarks associated with the implantation window, we performed integrated transcriptomic and proteomic profiling of WOI assembloids, with CTRL and SEC assembloids (Fig. 3A, Fig. S3A∼S3E).

**Fig 3.**
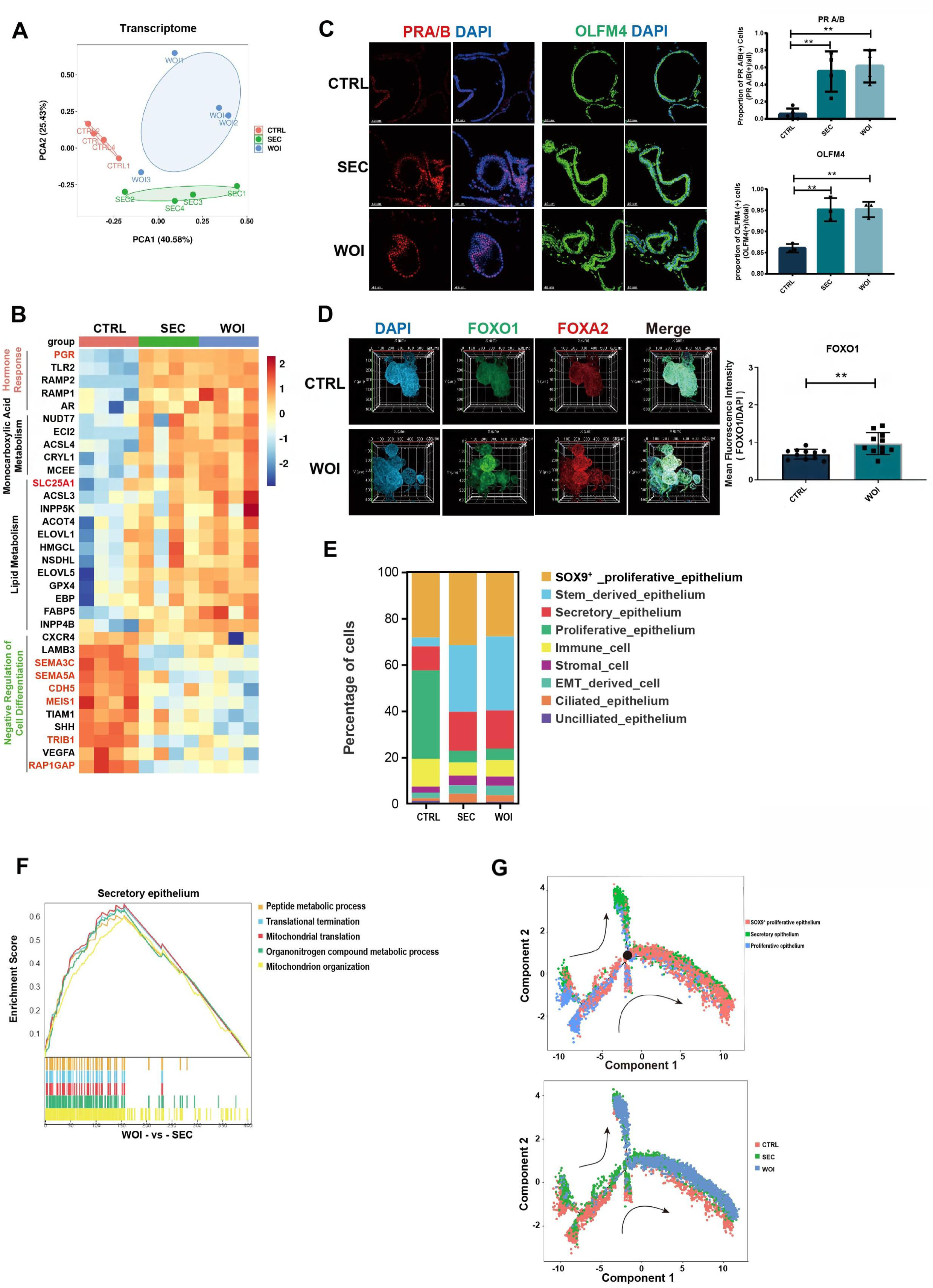
Receptive endometrial assembloids recapitulate WOI-associated hormone response (A) Principal component analysis (PCA) plot computed with differentially expressed genes in the bulk transcriptome of endometrial assembloids belonging to the CTRL, SEC and WOI groups. (B) Heatmap showing that enrichment of differentially expressed genes for the terms of hormone response, monocarboxylic acid metabolism, lipid metabolism, and negative regulation of cell differentiation. The color represents log-transformed fold change of gene expression. (C) Responsiveness to progesterone and estrogen was evaluated by IF to PRA/B and OLFM4 with IF respectively. Scale bar = 40 μm, **P ≤ 0.005. (D) Exhibition of implantation marker (FOXO1) and endometrial gland marker (FOXA2) through combination of assembloid clearing, IF and light sheet microscopy. Nuclei were counterstained with DAPI. **P ≤ 0.005. (E) Bar graph exhibiting various percentages of each cell type in the three groups. (F) GSEA between the SEC and WOI groups for secretory epithelium. (G) Pseudotime trajectory showing the transformation between proliferative and secretory epithelium in the CTRL, SEC and WOI groups. Arrows indicate the direction of the pseudotime trajectory. The black dot indicates the key differentiation node.

WOI assembloids exhibited a robust hormone response, as evidenced by upregulated PGR expression at the transcriptome level (Fig. 3B). Integrated multi-omics analysis identified 179 genes/proteins that were significantly upregulated in WOI versus CTRL assembloids, the majority of which are implicated in estrogen signaling pathways (Fig. S3F∼S3G). Furthermore, immunostaining for PRA/B, which was used to quantify progesterone responsiveness, revealed the highest signal intensity in the WOI group, concurrent with upregulation of the estrogen-responsive protein OLFM4 (Fig. 3C). FOXO1, a crucial marker of endometrial receptivity reliant on PGR signaling, was significantly elevated in WOI compared to CTRL assembloids (Fig.3D, Video S5). These observations suggest that progesterone signaling is central to the establishment of the WOI phenotype in assembloids, and collectively demonstrate that WOI assembloids mount a robust response to both estrogen and progesterone.

Hormonal stimulation induced an expansion of secretory epithelium and a concomitant reduction in proliferative epithelium in SEC and WOI assembloids, indicative of the proliferative-to-secretory phase transition (Fig. 3E). At the single-cell level, the secretory epithelium, which is critical for implantation window, was enriched for genes involved in cellular metabolic processes and hypoxic responses (HIF-1 signaling pathway) (Fig. S2G∼S2H). Compared to SEC assembloids, secretory epithelium in WOI assembloids exhibited enhanced peptide metabolism and mitochondrial energy metabolism (Fig. 3F), which are functional adaptations that support endometrial decidualization and embryo implantation. Single-cell trajectory analysis further revealed that proliferative epithelium differentiates into secretory epithelium under the regulation of key nodal genes defining the transition between states 5 and 6, including KRT19, MALAT1, MT2A, and the RPL and RPS families (Fig. 3G, Fig. S2J∼S2L). Notably, WOI assembloids displayed a more complete proliferative-to-secretory epithelial transition than SEC assembloids (Fig. 3G).

Overall, WOI assembloids possessed the hormone response characteristic of implantation window, closely resembling the gene traits of embryo implantation.

### Receptive endometrial assembloids possess enhanced energy metabolism

WOI assembloids exhibited upregulation of monocarboxylic acid and lipid metabolism (represented by SLC25A1(38)), and hypoxia response (represented by HIF1α(26)) (Fig.3B, Fig.S3G∼S3I). Likewise, the secretory epithelium, critical for the implantation window, contributed to cellular metabolic processes and HIF-1 signaling pathway response to hypoxia at the single-cell level (Fig. S2G∼S2H).

To further investigate this trend, the Mfuzz algorithm was utilized to analyze gene expression across these three groups, focusing on gene clusters that were progressively upregulated or downregulated. Mitochondrial-related genes were found to exhibit the highest expression levels in WOI endometrial assembloids (Fig. 4A). Concordantly, at the protein level, WOI endometrial assembloids maintained significantly higher expression of mitochondrial proteins compared to SEC assembloids (Fig. 4B). TEM analysis further revealed that WOI endometrial assembloids possessed the largest average mitochondrial area among the three groups (Fig. 4C). The expression of mitochondrial-related genes increased from CTRL to SEC to WOI assembloids, with COA1 ensuring proper nuclear-mitochondrial connection(39), OXA1L promoting mitochondrial translation(40), and TIMMDC1 being crucial for mitochondrial complex I assembly(41) (Fig. 4D). WOI assembloids notably expressed higher levels of OXA1L and TIMMDC1 than the SEC assembloids (Fig. 4D). Furthermore, WOI assembloids produced more ATP and IL8(42) (Fig. 4E).

**Fig 4.**
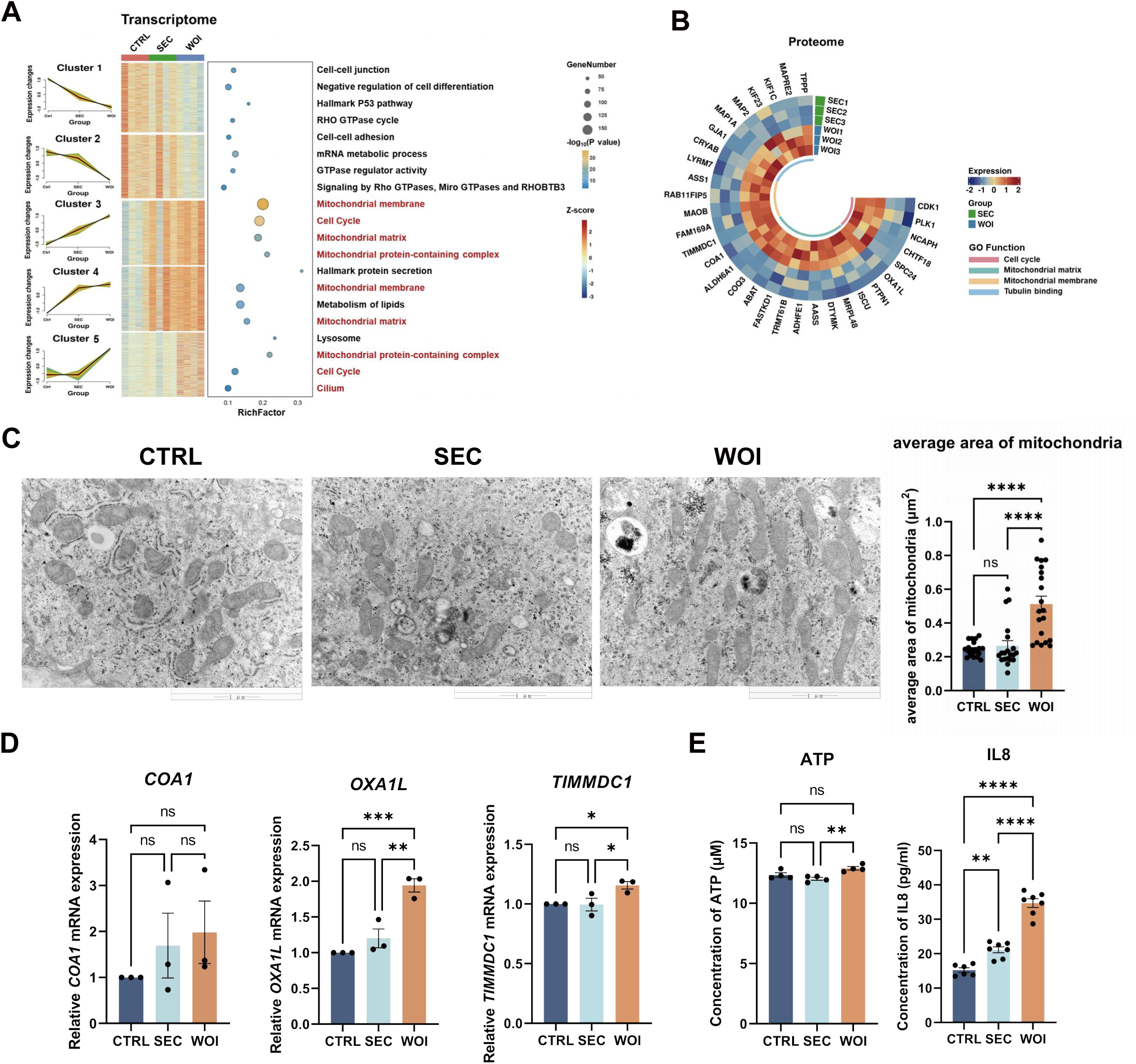
Receptive endometrial assembloids possess enhanced energy metabolism (A) The Mfuzz trend analysis displayed the transcriptional variation trends of five clusters from CTRL, SEC, and WOI groups (with a focus on the differences between SEC and WOI assembloids). The heatmap illustrated corresponding gene expression profile (where color represents Z-score). The bubble plot showed the associated GO functions (with bubble size representing the number of genes and bubble color indicating the P value). (B) The circular heatmap illustrated the functional differences of SEC and WOI assembloids at the proteomic level. The color represents protein expression levels, and the innermost circle color represents GO functions. (C) Transmission electron microscopy images displayed the mitochondrial morphology of CTRL, SEC, and WOI assembloids, along with a quantitative comparison of mitochondrial area. Scale bar = 1 μm. (D) RT-qPCR assessed the expression levels of mitochondrial function-related genes in the three assembloid groups. (E) Quantitative comparison of the concentration of ATP (left) and IL8 (right) released by CTRL, SEC and WOI assembloids. *P<0.05,**P<0.005,***P<0.0005,****P<0.0001.

Thus, compared to SEC assembloids, WOI assembloids exhibited enhanced energy metabolism, underpinned by larger, functionally competent mitochondria.

### Receptive endometrial assembloids increased the ciliary assembly and motility

Cilia are specialized structural components of the endometrium, whose growth and development, assembly and movement are essential for establishing the endometrial implantation window and facilitating embryo implantation. Under the electron microscopy, cilia were most observed in the WOI assembloids (Fig. 2E). Concordantly, cilia-related genes and proteins exhibited peak expression levels in WOI endometrial assembloids (Fig. 5A∼5B). Transcriptomic profiling further demonstrated that, relative to SEC assembloids, WOI assembloids displayed upregulated expression of genes linked to ciliary assembly, ciliary basal body, and motile cilia, whereas genes associated with non-motile cilia were downregulated (Fig. 5A). Key differentially expressed genes in WOI assembloids included *NEK2* (ciliary assembly), *CFAP36* (ciliary basal body), and *DNAH9* and *TPPP* (motile cilia), with NEK2 showing the most pronounced upregulation (Fig.5B). At the protein level, WOI assembloids also exhibited elevated expression of factors involved in ciliary assembly (TBC1D1, IFT22, IFT57), ciliary basal body biogenesis (PJA2), and motile cilia function (Fig. 5C).

**Fig 5.**
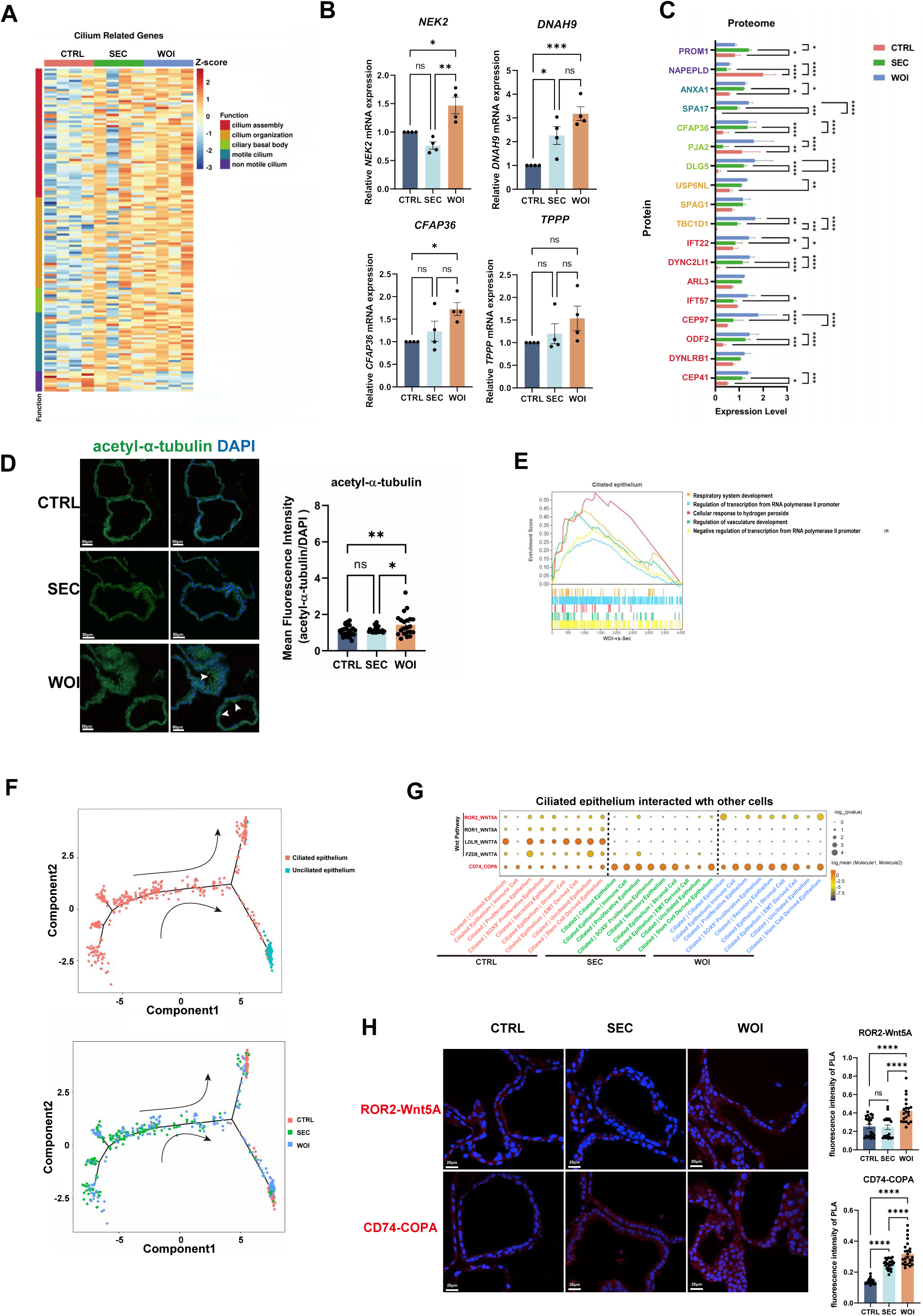
Receptive endometrial assembloids increased the ciliary assembly and motility (A) The heatmap illustrated the expression of cilia-related genes across the CTRL, SEC, and WOI assembloids. The color represents Z-score, while the leftmost block indicates various characteristic functions related to cilia. (B) RT-qPCR assessed the expression levels of cilia-related genes in the three assembloids. (C) The histogram showed the expression of cilia-related proteins in the three groups of assembloids. The color of the longitudinal protein names corresponds to the color of cilia-related functional blocks in Fig 5A. (D) IF analysis of cilia assembly marked by acetyl-α-tubulin. Nuclei were counterstained with DAPI. The arrowhead indicates cilia. Scale bar = 50 μm. (E) GSEA between the SEC and WOI groups for ciliated epithelium. (F) Pseudotime trajectory showing the transformation between ciliated and unciliated epithelium in the CTRL, SEC and WOI groups. Arrows indicate the direction of the pseudotime trajectory. (G) Dot plots demonstrating the Cellphone DB analysis of relevant receptors and ligands of ciliated epithelium with other cell types. The size of the dot represents the level of significance. The color of the dot indicates the mean of the average expression level of interacting molecule 1 in ciliated epithelium and molecule 2 in other cell types. (H) Proximity ligation assay (PLA) validating the interactions of ROR2-Wnt5A and CD74-COPA in the CTRL, SEC and WOI assembloids. Red signals the interaction of two proteins. Nuclei were counterstained with DAPI. Scale bar = 20 μm. *P<0.05,**P<0.005,***P<0.0005,****P<0.0001.

Additionally, acetyl-α-tubulin (cilia marker (15)) were highly expressed in the WOI assembloids (Fig. 5D).

Single-cell transcriptome analysis revealed hormone treatment increased ciliated epithelium and decreased unciliated epithelium in SEC and WOI groups (Fig. 3E). Ciliated epithelium functioned in protein binding, cilium organization and assembly, while unciliated epithelium acted on actin cytoskeleton and translation (Fig. S2E∼F). Notably, WOI assembloids’ ciliated epithelium additionally regulated vasculature development and displayed higher transcriptional activity than the SEC group (Fig. 5E). The ciliated to unciliated epithelial transition occurred during the menstrual cycle, which was regulated by key genes, such as *GAS5, JUN, RPL* and *RPS* families (Fig. 5F, Fig.S2M∼S2O). Given that these functional specializations are likely mediated by intercellular crosstalk, we employed CellPhoneDB (a computational tool for predicting ligand–receptor interactions (43)) to investigate cell–cell communication networks. Ciliated epithelium of WOI assembloids interacted with immune cells and secretory epithelium, showing enhanced invasion ability via CD74-COPA(44), and ROR2-WNT5A(45), which was validated by proximity ligation assay (PLA) (Fig. 5G∼5H).

In summary, the WOI assembloids revealed accumulated ciliated epithelium’s role in preparing the implantation window, with increased ciliary assembly and motility compared to SEC assembloids.

### Receptive endometrial assembloids experienced epithelial-mesenchymal transition (EMT)

The WOI assembloids displayed upregulated cell differentiation not only at assembloid level (Fig. 3B) but also at cellular level, which is represented by EMT. EMT is a common and crucial biological event in the endometrium during the implantation window(46). During the EMT process, epithelial cells lose their epithelial characteristics while gaining migratory and invasive properties of fibroblasts. Integrated transcriptomic and proteomic analyses revealed increased EMT in WOI assembloids (Fig. S3G).

EMT-derived cells, exhibiting gene expression patterns typical of epithelial and stromal cells, as well as EMT, are more abundant in the SEC and WOI groups, and act on protein binding, cell cycle, organelle organization, and reproduction (Fig. S2E). They performed enhanced lamellipodium-mediated cell migration, cell junction and cytoskeleton regulation in the WOI assembloids compared to the SEC assembloids (Fig. S4A). Single-cell trajectory analysis further revealed that WOI assembloids underwent a more complete differentiation from proliferative epithelium to EMT-derived cells than SEC assembloids, a process governed by key regulatory genes including *DOC2B, FXYD3,* and *LPCAT3* (Fig. S4B∼S4D). EMT-derived cells and epithelium coordinate functionally during the implantation window (Fig. S4E). Specifically, EMT-derived cells highly expressed NRP1 and SLC7A1, while their receptors (SEMA3A and CSF1) were more upregulated in the epithelium of WOI versus SEC assembloids (Fig. S4E, S4G). NRP1-SEMA3A has been reported to promote vascularization and responds to hypoxia(47). SLC7A1(48) and CSF1(49) both support receptivity establishment, embryo implantation and development. Compared with canonical stromal cells, EMT-derived cells exhibited distinct patterns of crosstalk with epithelial or immune cells. For instance, CD44 and CD46, known for their roles in cell adhesion (50) and immunoregulation(51), are highly expressed in stromal cells and EMT-derived cells, respectively, and bind separately with SPP1(in epithelial cells) and JAG1 (in stromal cells) (Fig. S4E∼S4G).

In general, WOI assembloids recapitulate the EMT-driven endometrial remodeling process that underpins the transition to the implantation window.

### The receptive endometrial assembloids possess the potential for embryo implantation

To validate WOI assembloids’ ability to recapitulate embryo implantation (a key biological process of receptive endometrium), we established an assembloid-based co-culture system. Given the rarity and ethical constraints associated with human embryos, we employed blastoids (corresponding to the human embryo at 6 days post-fertilization, referred to as “Day 6”) for implantation into the endometrial assembloids (Fig.6A). By Day 9, blastoids were observed to survive and integrate within the endometrial assembloids (Fig.6B). Importantly, co-cultured blastoids retained the capacity for normal tri-lineage differentiation, as evidenced by the appropriate expression of epiblast (OCT4), hypoblast (GATA6), and trophoblast (KRT18) lineage markers (Fig.6C). We next quantified blastoids survival rates and their rates of interaction with endometrial assembloids across the CTRL, SEC, and WOI groups. Strikingly, blastoids co-cultured with WOI assembloids exhibited significantly higher survival and interaction rates compared to CTRL and SEC assembloids, with survival rates of 66% (WOI), 19% (CTRL), and 28% (SEC), and interaction rates of 90% (WOI), 47% (CTRL), and 53% (SEC) respectively (Fig.6D∼E). This demonstrates that WOI assembloids possess a markedly enhanced capacity to support blastoids survival and integration, highlighting their functional relevance to the receptive endometrium.

**Fig 6.**
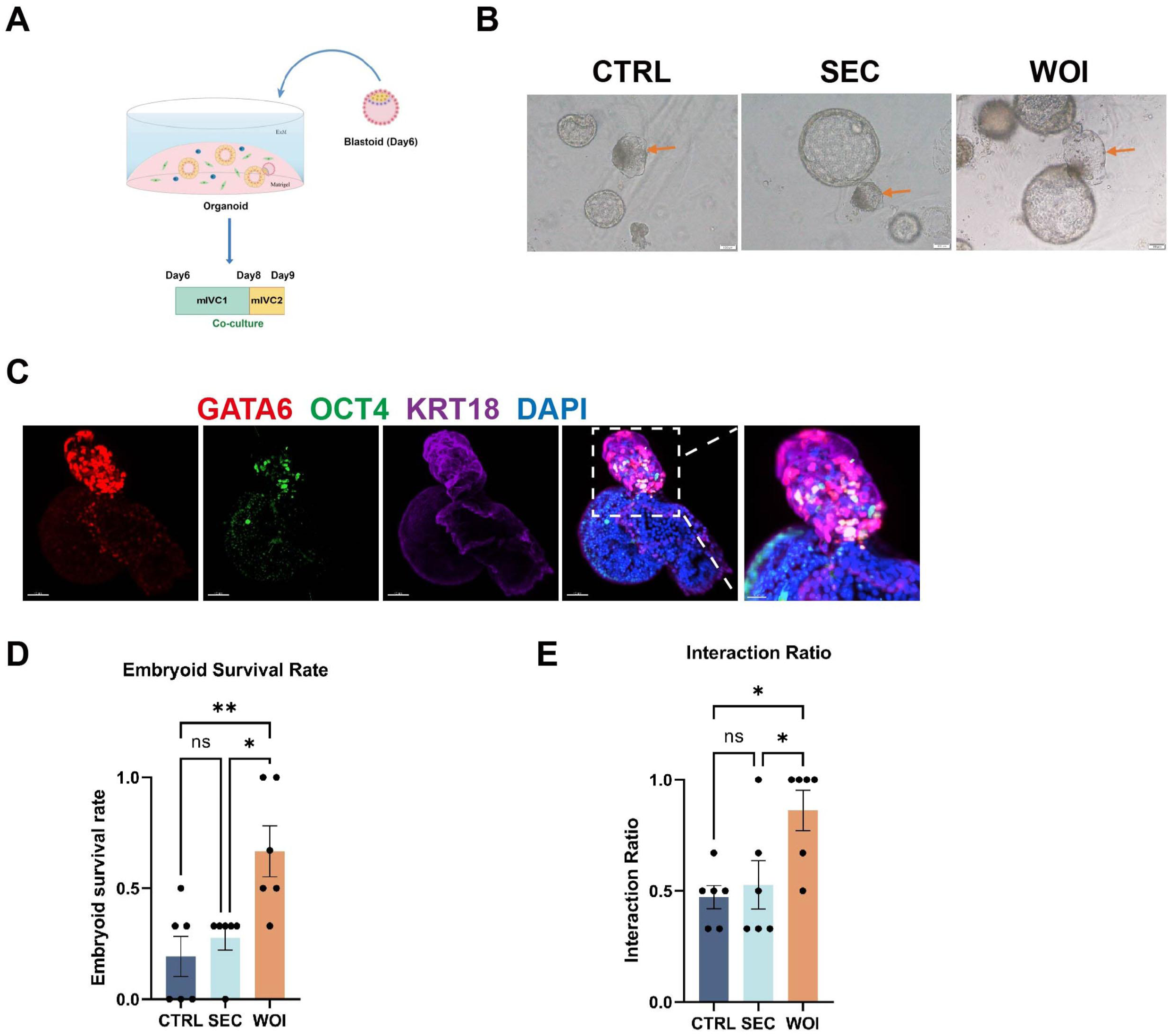
The receptive endometrial assembloids possess the potential for embryo implantation. (A) Diagram illustrated the co-culture model of endometrial assembloids with blastoids (the blastoid stage corresponds to a 6-day post-fertilization human embryo, referred to as Day 6 here). mIVC1: modified In Vitro Culture Medium 1, mIVC2: modified In Vitro Culture Medium 2. (B) Bright-field images of the co-culture of CTRL, SEC, and WOI assembloids with blastoids (Day 9) (yellow arrows indicate the blastoids). Scale bar = 100μm. (C) Whole-mount fluorescence staining of Day 9 co-cultured embryoid and assembloid. OCT4 indicates the epiblast, GATA6 indicates the hypoblast, and KRT18 indicates the trophoblast. Scale bar = 40μm and 20μm (the rightmost image). (D) Comparison of the survival rates of Day 9 embryoid in CTRL, SEC, and WOI assembloids. (E) Comparison of the interaction ratios between Day 9 embryoid and endometrial assembloids in the CTRL, SEC, and WOI groups. *P < 0.05, **P < 0.005.

In summary, WOI endometrial assembloids not only exhibited the typical structural and molecular features of the implantation window but also demonstrated significant potential for embryo implantation.

## Discussion

In this study, we constructed the WOI endometrial assembloids, and observed the remarkable resemblance in structure and function to the in vivo endometrium. The assembloids consist of three primary types of endometrial cells, specifically epithelial, stromal, and immune cells, with epithelial cells assembling into glands surrounded by immune cells and stromal cells. This conservation of cellular composition and tissue architecture provides a foundational basis for mimicking the receptive endometrium.

While previous studies have induced secretory phase transition in endometrial assembloids using E2, P4, and cAMP, methodologies for accurately simulating the WOI or mid-secretory endometrium remain inadequate and require further refinement. Prior research has indicated that placental signals can enhance endometrial assembloid differentiation (2), with PRL, hCG and HPL implicated in decidualization, implantation, immunoregulation, and angiogenesis (21, 22, 24). Specifically, PRL, synthesized by the adenohypophysis, endometrium, and myometrium, plays a vital role in implantation, immunoregulation, and angiogenesis, with the secretory endometrium producing PRL in response to MPA and E2, leading to ciliated cell formation and stromal cell decidualization (2, 21). HCG, secreted by trophoblasts in early pregnancy, modulates decidual cells (22) and improves endometrial thickness and receptivity(23). The introduction of hCG has been shown to upregulate key receptivity-associated factors, such as endocytosis proteins, hypoxia-inducible factor 1 (HIF1), chemokines, and glycodelin (52). HPL contributes to the development and function of uterine glands(24). Consequently, supplementary PRL, hCG and HPL in our system augmented hormone responsiveness and receptivity, promoted cellular differentiation, and ultimately generated a model more representative of the receptive endometrium.

Notably, WOI assembloids exhibited the characteristic ultrastructures, such as cilia. Motile cilia are present in endometrial epithelium throughout the human menstrual cycle (1). These hair-like organelles extend from the cell surface and beat rhythmically, facilitating cell and tissue movement while driving fluid transport across the epithelium (53). During endometrial decidualization, the number and length of cilia increase, a process driven by the two primary regulatory factors for embryo implantation: estrogen and progesterone (15, 54, 55). Throughout the menstrual cycle, ciliated and unciliated epithelia undergo mutual transformation from the secretory phase (or mid-secretory phase) to the menstrual phase, and then to the proliferative phase. Ciliated cell abundance peaks in the early-to-mid-secretory endometrium (1), with the mid-secretory phase exhibiting a higher proportion of ciliated cells as opposed to the early and late-secretory phases (25). However, with aging, the expression of cilia-related genes in the endometrium is downregulated(56). Recurrent implantation failure (RIF) patients often present with endometrial ciliary defects, leading to impaired decidualization and repeated implantation failure (55). Thus, cilia play a crucial role in endometrial decidualization and embryo implantation. Consistently, the WOI assembloid exhibited increased ciliated epithelium, along with enhanced assembly and motility of cilia.

Furthermore, WOI assembloids demonstrated energy and lipid metabolism patterns resembling those of the in vivo receptive endometrium. Enhanced energy metabolism involves monocarboxylic acid metabolism and mitochondrial oxidative phosphorylation. Monocarboxylic acids, exemplified by lactate, pyruvate, and ketone bodies, are vital metabolites in most mammalian cells. During the implantation window, elevated lactate levels mobilize endometrial monocarboxylic acid metabolism and function as a pregnancy-related signal, stimulating secretion in the epithelium. Subsequently, ATP, the primary product of energy metabolism, induces neighboring epithelial cells to release IL8, promoting decidualization of stromal cells (42). Concordantly, our WOI assembloids indeed possessed larger and functionally active mitochondria, and produced much more ATP and IL8 than CTRL and SEC assembloids. Lipid metabolism, responsible for energy storage, signal transduction, cell proliferation, apoptosis, and membrane trafficking, plays a crucial role in endometrial receptivity and implantation, although its precise mechanisms remain incompletely understood (57–59). Thus, WOI assembloids possessed metabolic characteristics of in vivo implantation window.

However, our WOI endometrial assembloids also exhibit some limitations. It is undeniable that the assembloids cannot perfectly replicate the in vivo endometrium, which comprises functional and basal layers with a greater abundance of cell subtypes, under superior regulation by hypothalamic-pituitary-ovarian (HPO) axis. Specifically, stromal and immune cells are challenging to stably passage, and their proportion is lower than in the in vivo endometrium. While the in vivo peri-implantation period exhibits intricate gene expression dynamics driven by systemic regulation, our models only partially recapitulate these changes, primarily in mitochondria-and cilia-associated genes. Nevertheless, to some extent, these WOI assembloids possess receptivity characteristics and can be utilized for investigating receptivity-related scientific questions or conducting in vitro drug screening. Further refinements are required to fully simulate the dynamic endometrial gene expression patterns across all menstrual cycle stages. We are looking forward to integrating stem cell induction, 3D printing, and microfluidic systems to modify the culture environment.

In summary, we developed a receptive endometrial assembloid that mimics in vivo implantation-window endometrium (Fig. 7). This model exhibits WOI-characteristic ultrastructures, including pinopodes and cilia, displays hormonal responsiveness and characteristic glycogen-secreting functions, recapitulates the processes such as decidualization, metabolic changes and EMT, and retains embryo implantation potential. It serves as a valuable platform to investigate peri-implantation endometrial physiology and pathology, maternal-fetal interactions, with potential practical applications and clinical translation.

**Fig. 7.**
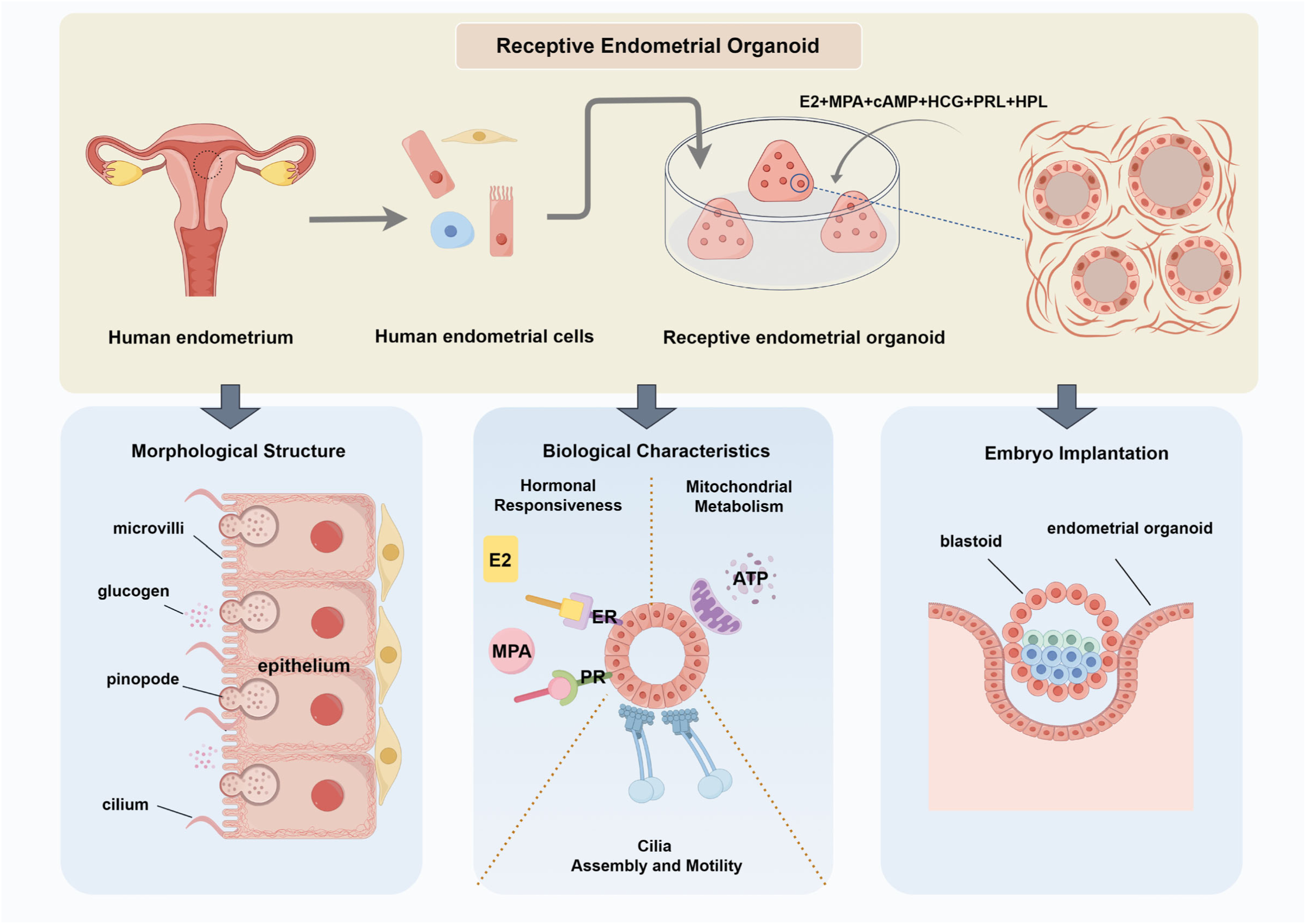
Schematic diagram displaying the establishment and validation of receptive endometrial assembloids, and summarizing the characteristic biological events of implantation-window endometrium.

## Materials and Methods

### Establishment of endometrial assembloids

All experiments involving human subjects followed medical ethical principles and the Declaration of Helsinki, and were approved by the Ethics Committee of Qilu Hospital of Shandong University. Informed consents were obtained from patients. Females of reproductive age who received hysterectomy for benign diseases were selected for this study. Clinical information of patients providing endometrial tissue was listed in Dataset S1. Experimental procedures are presented in detail in the Supporting Information.

### Hormone treatment of endometrial assembloids

Hormonal treatment was initiated following the assembly of the endometrial assembloids (about 7-day growth period). The hormone regimen for inducing endometrial secretory phase, as described by Thomas E. Spencer et al (9), consisted of estradiol for 2 days followed by a combination of estradiol (Sigma E2758), medroxyprogesterone acetate (Selleck S2567) and N6,2′-O-dibutyryladenosine 3′,5′-cyclic monophosphate sodium salt (cAMP) (Sigma D0627) for 6 days.

To simulate the endometrium of implantation window, we developed a model by incorporating various pregnancy-related hormones. The hormone regimen included estradiol for the first two days, followed by a combination of estradiol, medroxyprogesterone acetate, cAMP, Human Chorionic Gonadotropin (HCG) (Livzon Pharmaceutical Group Inc), Human Placental Lactogen (HPL) (R&D Systems 5757-PL), and prolactin (Peprotech 100-07) for 6 days. The CTRL group was cultured in ExM without hormone supplementation and subjected to parallel culture for 8 days along with the two aforementioned groups. (Fig. 1A) (Table S2)

## Supporting information

Video_Table_Dateset

## ACKNOWLEDGMENTS

This work was supported by National Natural Science Foundation of China (82192874 to H.Z), the National Key Research and Development Program of China (2021YFC2700301 to K.W), the Basic Science Center Program (31988101 to Z-J.C.), Fundamental Research Funds for the Central Universities(2022JC006 to K.W), the Fundamental Research Funds of Shandong University Taishan Scholars Program of Shandong Province(ts20190988 to H.Z.; tsqn201909194 to K.W), Innovative research team of high-level local universities in Shanghai (SHSMU-ZLCX20210201 to K.W).

We sincerely appreciate Prof. Tianqing Li (Kunming University of Science and Technology) and Prof. Shaorong Gao (Tongji University) for insightful comments and discussions.

We are grateful to Guangzhou Genedenovo Biotechnology Co., Ltd for assisting in sequencing and bioinformatics analysis. We also thank to YiKon Medical from China for technological assistance of ERA, Jingjie PTM Biolab (HangZhou) Co., Inc. for proteomics.

## Author contributions

H. Zhao, K. Wu and Z.-J.C conceived the project and supervised research. Y. Zhang, R. Zhao and H. Zhao designed the experiments. Y. Zhang performed the experiments, analyzed and interpreted data. Y. Zhang wrote the manuscript. R. Zhao, Y. Li, B. Liu, and H. Zhao revised the manuscript. P. Liu provided human endometrium samples. C.Yang, J. Song, T. Li, C. Yin, M. Lu, Z. Hou and C. Zhang provided technical support. All authors revised and approved the manuscript.

The authors declare no competing interest.

---

**Fig. S1.**
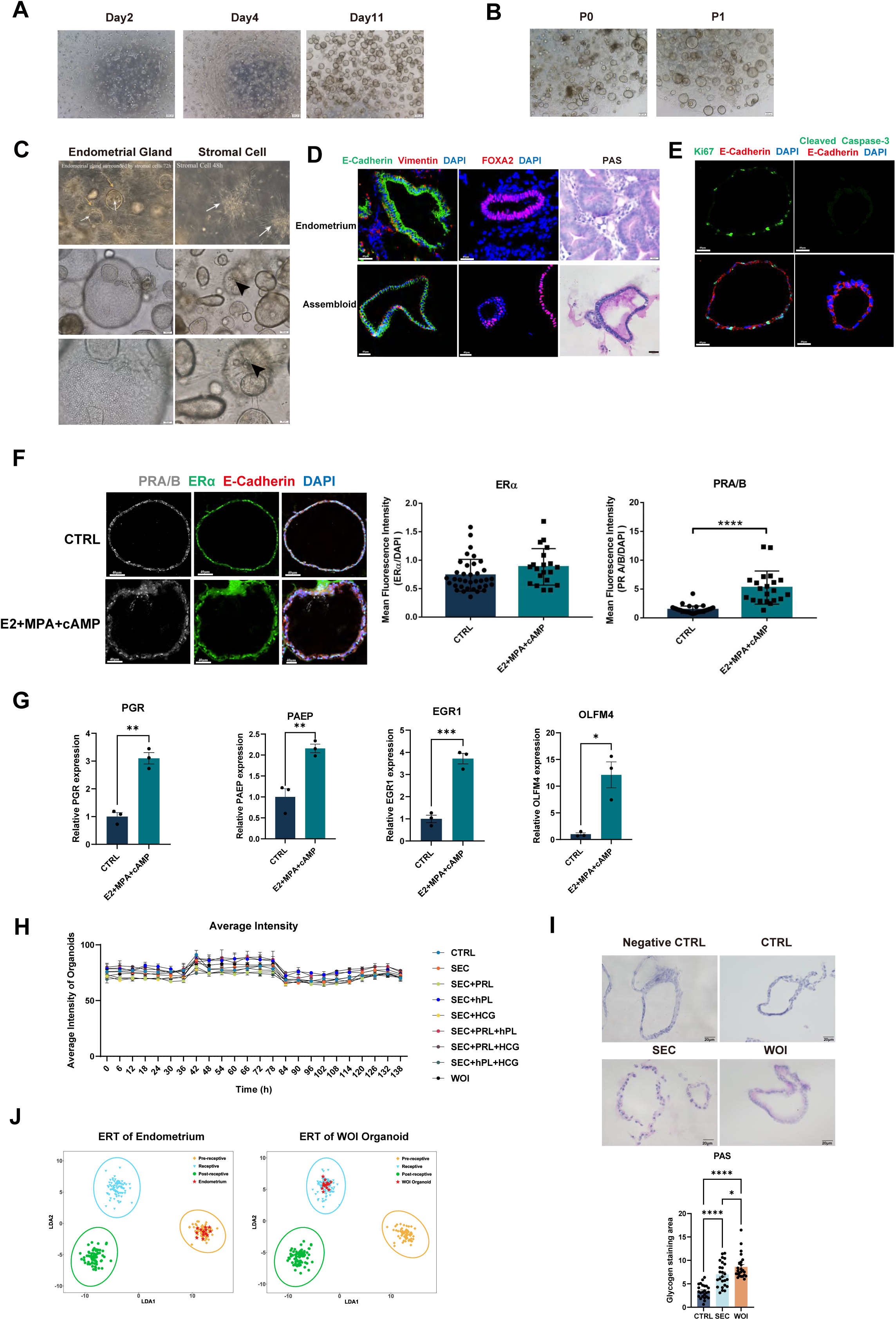
Developing receptive endometrial assembloids in vitro (A) Brightfield of endometrial assembloids on day2, day4 and day11. Scale bar = 200 μm. (B) Brightfield of endometrial assembloids in the primary generation (P0) and first generation (P1). Scale bar = 200 μm. (C) Screenshot of video S1 showing endometrial glands gradually developing into a vesicular shape, and the surrounding stromal cells arranging in a fibrous pattern in the CTRL endometrial assembloids (100x, up-left) (The yellow arrows indicate stromal cells and the white arrows indicate endometrial glands). Screenshot of video S2 displaying the stromal cells growing in fibrous pattern and forming an extensive network in the CTRL endometrial assembloids (200x, up-right) (The white arrows indicate stromal cells). The epithelial cells arrange like paving stones (middle and down, left). Stromal cells formed an extensive network (middle and down, right) (The arrowhead indicates stromal cells). Scale bar = 100 μm (middle), Scale bar = 50 μm (down). (D) Validation of epithelial, stromal cell and endometrial gland markers (E-cadherin, vimentin and FOXA2, respectively) with immunofluorescence (IF) in the endometrium in vivo and CTRL endometrial assembloids in vitro. Nuclei were counterstained with DAPI. Scale bar = 40 μm. Periodic acid-schiff staining (PAS) of endometrium in vivo and endometrial assembloid in vitro. Scale bar = 20 μm. (E) IF analysis of proliferation and apoptosis indicated by Ki67 and cleaved caspase-3 in the CTRL assembloids, respectively. Nuclei were counterstained with DAPI. Scale bar = 40 μm. (F) Verification of hormone responsiveness by the expression levels of ERα and PRA/B (E-cadherin indicated the marker of epithelial cell). Scale bar = 40 μm. ***P ≤ 0.0005. (G) Relative expression of PGR, PAEP, EGR1, and OLFM4 in the CTRL and hormone-treated assembloids by RT-qPCR. *P ≤ 0.05, **P ≤ 0.005, ***P ≤ 0.0005. (H) The dynamic changes of the average intensity of assembloids over time in each hormone regimen. (I) PAS of CTRL, SEC and WOI assembloids. PAS staining with diastase digestion was Negative CTRL. Scale bar = 20 μm. Quantitative comparison of glycogen staining area in the CTRL, SEC and WOI assembloids. *P ≤ 0.05, ****P ≤ 0.0001. (J) Endometrial receptivity evaluation of endometrium and their derived WOI assembloids through ERA. Asterisks indicate individual samples.

**Fig S2.**
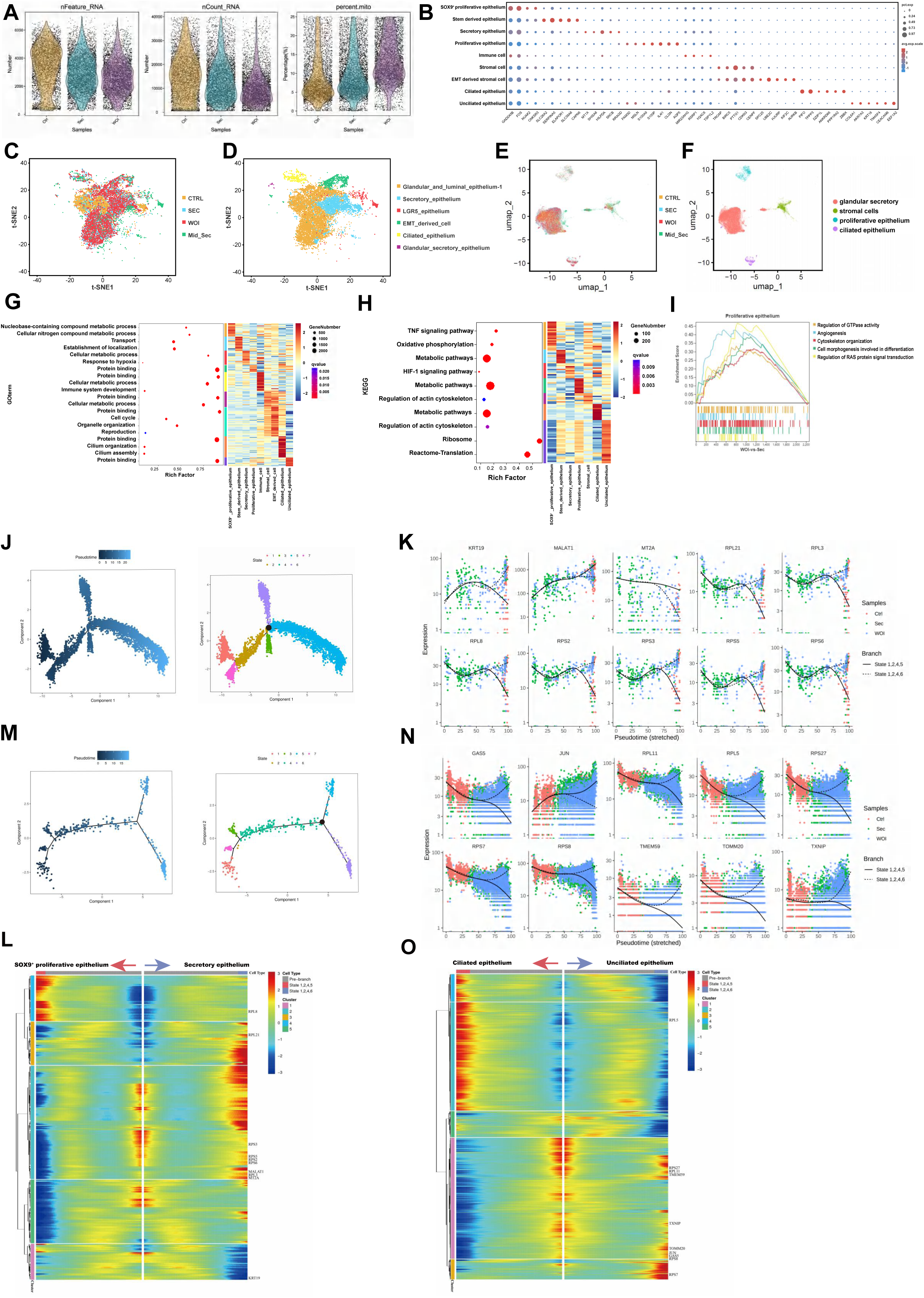
Various functions performed by all kinds of cells identified with scRNA-seq. (A) Box plot of the gene numbers detected, UMI numbers, and ratios of mitochondrial gene expression in single cells of the CTRL, SEC and WOI groups. (B) Bubble diagram showing the distribution of marker gene expression in each cluster. (C∼D) Comparison of cell composition between CTRL, SEC, WOI endometrial assembloids and mid-secretory endometrium demonstrating samples (Stephen R. Quake in 2020) (C) and cell types (D). (E∼F) Comparison of cell composition between CTRL, SEC, WOI endometrial assembloids and mid-secretory endometrium demonstrating samples (Garcia-Alonso in 2021) (E) and cell types (F). (G) Bubble diagram and heatmap showing the corresponding upregulated genes and GO function of each cluster of endometrial cells. (H) Bubble diagram and heatmap showing corresponding upregulated genes and KEGG functions of SOX9^+^ proliferative epithelium, stem-derived epithelium, secretory epithelium, proliferative epithelium, stromal cells, ciliated epithelium and unciliated epithelium. Color is proportional to log-transformed fold change of gene expression. (I) GSEA between the SEC and WOI groups for proliferative epithelium. (J) The single-cell pseudotime trajectory of SOX9^+^ proliferative epithelium, secretory epithelium and proliferative epithelium. Cells start at proliferative epithelium and progress to SOX9^+^ proliferative epithelium and secretory epithelium (left). There are seven major states over pseudotime (right). The black spot indicates the differentiation node between state 5 and state 6, indicating the direction from proliferative epithelium to SOX9^+^ proliferative epithelium and secretory epithelium, respectively. (K)(N) The horizontal axis is the pseudotime point, and the vertical axis is the gene expression level. The solid line represents states 1, 2, 4, and 5 corresponding to Fig. S2L (K) or Fig. S2O (N). Different colors represent samples in the CTRL, SEC and WOI groups. (L)(O) Heatmap of genes at the branch node regulating differentiation into SOX9^+^ proliferative epithelium or secretory epithelium (L), and ciliated epithelium or unciliated epithelium (O). The horizontal axis is the pseudo-time point (the pseudo-time point gradually increases from the middle to both sides). The vertical axis is the gene expression level, representing two differential directions on the left and right sides. Clusters represent the gene sets with a similar branch gene expression trend. Different colors represent the level of gene expression. (M) Pseudotime trajectory of ciliated and unciliated epithelium. Cells start at ciliated epithelium and progress to unciliated epithelium (left). There are seven major states over pseudotime (right). The black spot indicates the differentiation node between state 5 and state 6, indicating the direction of ciliated and unciliated epithelium, respectively. Arrows indicate the direction of the pseudotime trajectory.

**Fig S3.**
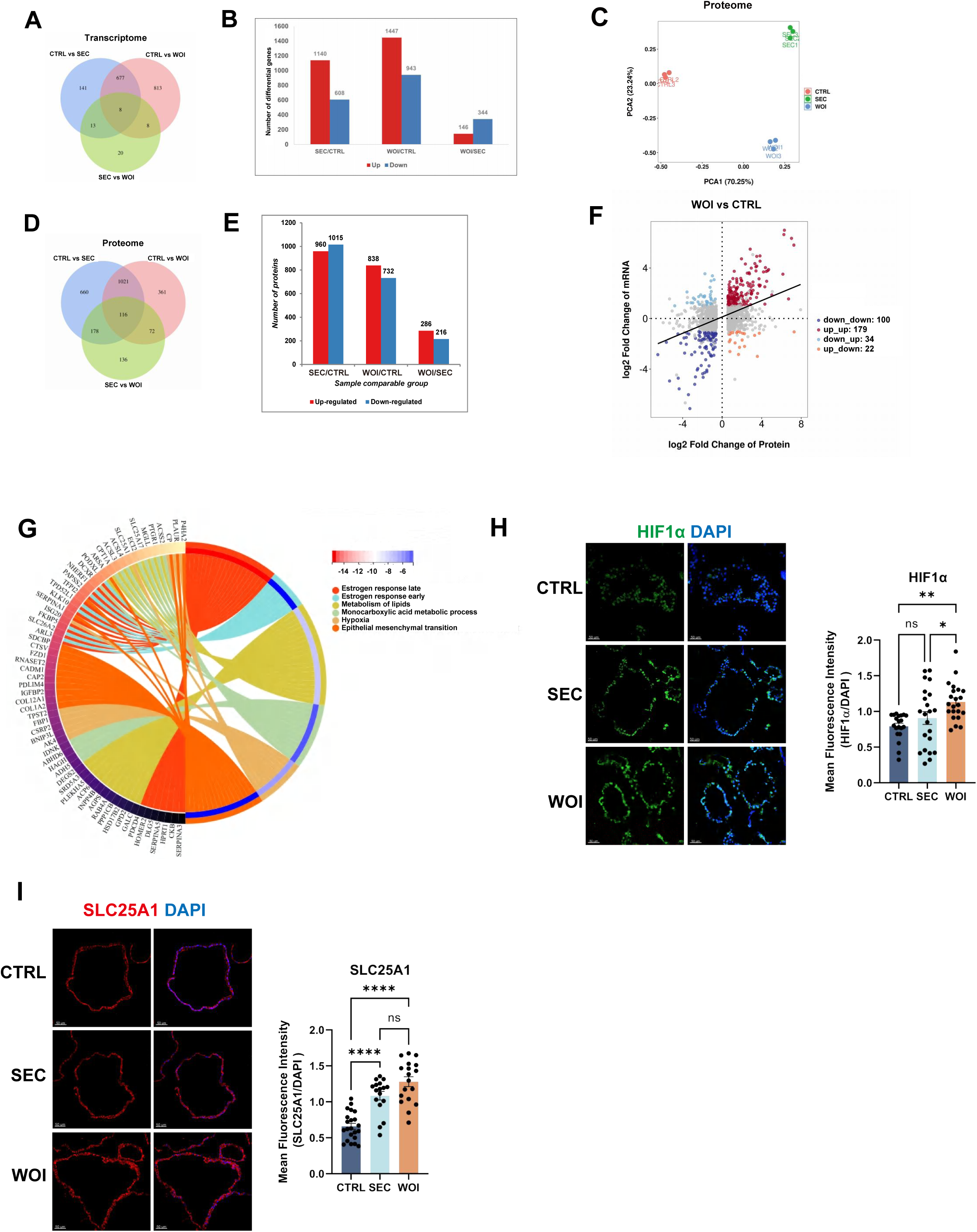
Comparisons between CTRL, SEC and WOI assembloids at the level of transcriptome and proteome. (A) Venn diagram showing differential genes in pairs among the three groups. Log_2_ FC (Fold Change)>1.2 or<-1.2, q value<0.05. (B) Histogram showing pairwise comparison of differential gene expression (DEG) from bulk RNA-seq among the CTRL, SEC and WOI groups. qvalue<0.05, |logFC|>1. (C) PCA plot computed with differentially expressed proteins in the micro proteomics of endometrial assembloids belonging to the CTRL, SEC and WOI groups. (D) Venn diagram showing differential proteins in pairs among the three groups. Log_2_ FC>1.2 or <-1.2, q value<0.05. (E) Histogram showing pairwise comparison of differential proteins in micro proteomics among the CTRL, SEC and WOI groups. (F) Scatterplot depicting the correlation between the transcriptome and proteome as for the WOI group and CTRL group. For transcriptome, FC>2 and p value<0.05 were defined as significantly upregulated. For proteome, FC>1.2 and p value<0.05 were defined as significantly upregulated. (G) Circle diagram showing the functions of genes upregulated in both the transcriptome and proteome as for the WOI group compared to CTRL group. (H) Verification of hypoxia response by the expression levels of HIF1α. Scale bar = 50 μm. *P ≤ 0.05, **P ≤ 0.005. (I) Verification of lipid metabolism by the expression levels of SLC25A1. Scale bar = 50 μm. ****P ≤ 0.0001.

**Fig S4.**
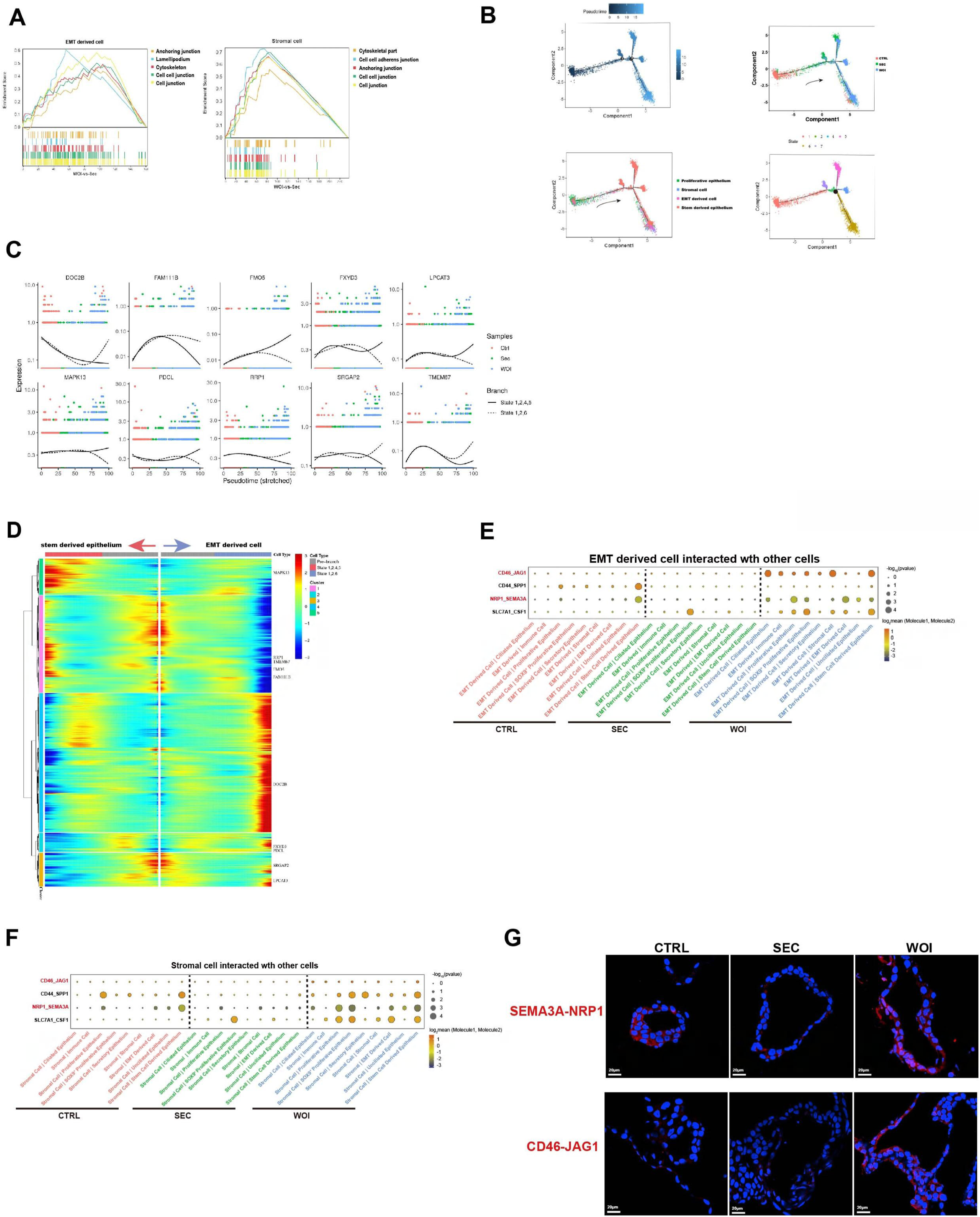
Receptive endometrial assembloids experienced epithelial-mesenchymal transition (EMT) (A) GSEA between the SEC and WOI groups for proliferative epithelium. (B) The single-cell pseudotime trajectory of proliferative epithelium, stromal cell, EMT derived cell and stem derived epithelium. Cells start at proliferative epithelium and progress to EMT derived cell. There are seven major states over pseudotime. The black spot indicates the differentiation node between state 4, 5 and state 6, indicating the direction from proliferative epithelium to EMT derived cell. Arrows indicate the direction of the pseudotime trajectory. (C) The horizontal axis is the pseudotime point, and the vertical axis is the gene expression level. The solid line represents states 1, 2, 4, and 5 corresponding to Fig. S4B. Different colors represent samples in the CTRL, SEC and WOI groups. (D) Heatmap of genes at the branch node regulating differentiation into EMT derived cell and stem-derived epithelium. The horizontal axis is the pseudo-time point (the pseudo-time point gradually increases from the middle to both sides). The vertical axis is the gene expression level, representing two differential directions on the left and right sides. Clusters represent the gene sets with a similar branch gene expression trend. Different colors represent the level of gene expression. (E-F) Dot plots demonstrating the Cellphone DB analysis of relevant receptors and ligands of EMT derived cell (E) or stromal cell (F) with other cell types. The size of the dot represents the level of significance. The color of the dot indicates the mean of the average expression level of interacting molecule 1 in EMT derived cells (E) or stromal cells (F) and molecule 2 in other cell types. (G) Proximity ligation assay (PLA) validating the interactions of SEMA3A-NRP1 and CD46-JAG1 in the CTRL, SEC and WOI assembloids. Red signals the interaction of two proteins. Nuclei were counterstained with DAPI. Scale bar = 20 μm.

## References

1. Wang W, et al. (2020) Single-cell transcriptomic atlas of the human endometrium during the menstrual cycle. Nat Med 26(10):1644–1653.

2. Turco MY, et al. (2017) Long-term, hormone-responsive organoid cultures of human endometrium in a chemically defined medium. Nat Cell Biol 19(5):568–577.

3. Boretto M, et al. (2017) Development of organoids from mouse and human endometrium showing endometrial epithelium physiology and long-term expandability. Development 144(10):1775–1786.

4. Cindrova-Davies T, et al. (2021) Menstrual flow as a non-invasive source of endometrial organoids. Commun Biol 4(1):651.

5. Shun Shibata SE, Luis A. E. Nagai, Eri H. Kobayashi, Akira Oike, Norio Kobayashi, Akane Kitamura, Takeshi Hori, Yuji Nashimoto, Ryuichiro Nakato, Hirotaka Hamada, Hirokazu Kaji, Chie Kikutake, Mikita Suyama, Masatoshi Saito, & Nobuo Yaegashi HO, Takahiro Arima (2024) Modeling embryo-endometrial interface recapitulating human embryo implantation. Sci Adv 10(8):eadi4819.

6. Cheung VC, et al. (2021) Pluripotent stem cell-derived endometrial stromal fibroblasts in a cyclic, hormone-responsive, coculture model of human decidua. Cell Rep 35(7):109138.

7. Gong L, et al. (2022) Bi-potential hPSC-derived Mullerian duct-like cells for full-thickness and functional endometrium regeneration. NPJ Regen Med 7(1):68.

8. Dolat L & Valdivia RH (2021) An endometrial organoid model of interactions between Chlamydia and epithelial and immune cells. J Cell Sci 134(5).

9. Harriet C. Fitzgerald PD, Susanta K. Behura, Danny J. Schust, and Thomas E. Spencer (2019) Self-renewing endometrial epithelial organoids of the human uterus. Proc Natl Acad Sci U S A (116(46)):23132–23142.

10. Wiwatpanit T, et al. (2020) Scaffold-Free Endometrial Organoids Respond to Excess Androgens Associated With Polycystic Ovarian Syndrome. J Clin Endocrinol Metab 105(3):769–780.

11. Thomas M Rawlings KM, Deborah M Taylor, Matteo A Molè, Katherine J Fishwick, Maria Tryfonos, Joshua Odendaal, Amelia Hawkes, Magdalena Zernicka-Goetz, Geraldine M Hartshorne, Jan J Brosens, Emma S Lucas (2021) Modelling the impact of decidual senescence on embryo implantation in human endometrial assembloids. eLife 2021;10:e69603.

12. Park SR, et al. (2021) 3D stem cell-laden artificial endometrium: successful endometrial regeneration and pregnancy. Biofabrication 13(4).

13. Park SR, et al. (2023) Development of cell-laden multimodular Lego-like customizable endometrial tissue assembly for successful tissue regeneration. Biomater Res 27(1):33.

14. Boretto M, et al. (2019) Patient-derived organoids from endometrial disease capture clinical heterogeneity and are amenable to drug screening. Nat Cell Biol 21(8):1041–1051.

15. Haider S, et al. (2019) Estrogen Signaling Drives Ciliogenesis in Human Endometrial Organoids. Endocrinology 160(10):2282–2297.

16. Cochrane DR, et al. (2020) Single cell transcriptomes of normal endometrial derived organoids uncover novel cell type markers and cryptic differentiation of primary tumours. J Pathol 252(2):201–214.

17. Bui BN, et al. (2020) Organoids can be established reliably from cryopreserved biopsy catheter-derived endometrial tissue of infertile women. Reprod Biomed Online 41(3):465–473.

18. Fitzgerald HC, Kelleher AM, Ranjit C, Schust DJ, & Spencer TE (2023) Basolateral secretions of human endometrial epithelial organoids impact stromal cell decidualization. Mol Hum Reprod 29(4).

19. Kirkwood PM, et al. (2022) Single-cell RNA sequencing and lineage tracing confirm mesenchyme to epithelial transformation (MET) contributes to repair of the endometrium at menstruation. Elife 11.

20. Paule SG, et al. (2021) Podocalyxin is a key negative regulator of human endometrial epithelial receptivity for embryo implantation. Hum Reprod 36(5):1353–1366.

21. Auriemma RS, et al. (2020) The Interplay Between Prolactin and Reproductive System: Focus on Uterine Pathophysiology. Front Endocrinol (Lausanne) 11:594370.

22. d’Hauterive SP, et al. (2022) Human Chorionic Gonadotropin and Early Embryogenesis: Review. Int J Mol Sci 23(3).

23. Nwabuobi C, et al. (2017) hCG: Biological Functions and Clinical Applications. Int J Mol Sci 18(10).

24. Bazer FW, Burghardt RC, Johnson GA, Spencer TE, & Wu G (2018) Mechanisms for the establishment and maintenance of pregnancy: synergies from scientific collaborations. Biol Reprod 99(1):225–241.

25. Garcia-Alonso L, et al. (2021) Mapping the temporal and spatial dynamics of the human endometrium in vivo and in vitro. Nat Genet 53(12):1698–1711.

26. Rocío Martínez-Aguilar LEK, Jane J Reavey, Hilary O D Critchley and Jacqueline A Maybin (2021) HYPOXIA AND REPRODUCTIVE HEALTH: The presence and role of hypoxia in the endometrium. Reproduction (161(1)):F1-F17.

27. Fatmous M, Rai A, Poh QH, Salamonsen LA, & Greening DW (2022) Endometrial small extracellular vesicles regulate human trophectodermal cell invasion by reprogramming the phosphoproteome landscape. Front Cell Dev Biol 10:1078096.

28. Jiang X, et al. (2022) Lack of VMP1 impairs hepatic lipoprotein secretion and promotes non-alcoholic steatohepatitis. J Hepatol 77(3):619–631.

29. Kuo WT, et al. (2021) The Tight Junction Protein ZO-1 Is Dispensable for Barrier Function but Critical for Effective Mucosal Repair. Gastroenterology 161(6):1924–1939.

30. Bissonnette L, et al. (2016) Human S100A10 plays a crucial role in the acquisition of the endometrial receptivity phenotype. Cell Adh Migr 10(3):282–298.

31. Srivastava C, et al. (2018) FAT1 modulates EMT and stemness genes expression in hypoxic glioblastoma. Int J Cancer 142(4):805–812.

32. Liu L, et al. (2020) si-SNHG5-FOXF2 inhibits TGF-beta1-induced fibrosis in human primary endometrial stromal cells by the Wnt/beta-catenin signalling pathway. Stem Cell Res Ther 11(1):479.

33. Geng J, Cui C, Yin Y, Zhao Y, & Zhang C (2022) LncRNA NEAT1 affects endometrial receptivity by regulating HOXA10 promoter activity. Cell Cycle 21(18):1932–1944.

34. Vargas MF, et al. (2012) Effect of single post-ovulatory administration of levonorgestrel on gene expression profile during the receptive period of the human endometrium. J Mol Endocrinol 48(1):25–36.

35. Kinnear S, Salamonsen LA, Francois M, Harley V, & Evans J (2019) Uterine SOX17: a key player in human endometrial receptivity and embryo implantation. Sci Rep 9(1):15495.

36. Huang P, et al. (2021) SOX4 facilitates PGR protein stability and FOXO1 expression conducive for human endometrial decidualization Elife 2022 Mar 4;11:e72073.

37. He A, et al. (2021) The role of transcriptomic biomarkers of endometrial receptivity in personalized embryo transfer for patients with repeated implantation failure. J Transl Med 19(1):176.

38. Yang Y, et al. (2021) SLC25A1 promotes tumor growth and survival by reprogramming energy metabolism in colorectal cancer. Cell Death Dis 12(12):1108.

39. Wang C, et al. (2019) MITRAC15/COA1 promotes mitochondrial translation in a ND2 ribosome–nascent chain complex. EMBO reports 21(1).

40. Itoh Y, et al. (2021) Mechanism of membrane-tethered mitochondrial protein synthesis. Science 371(6531):846–849.

41. Wang T, et al. (2021) C9orf72 regulates energy homeostasis by stabilizing mitochondrial complex I assembly. Cell Metabolism 33(3):531–546.e539.

42. Gu XW, et al. (2020) ATP mediates the interaction between human blastocyst and endometrium. Cell Prolif 53(2):e12737.

43. Efremova M, Vento-Tormo M, Teichmann SA, & Vento-Tormo R (2020) CellPhoneDB: inferring cell-cell communication from combined expression of multi-subunit ligand-receptor complexes. Nat Protoc 15(4):1484–1506.

44. Li M, et al. (2022) BUB1 Is Identified as a Potential Therapeutic Target for Pancreatic Cancer Treatment. Front Public Health 10:900853.

45. Li T, et al. (2022) WNT5A Interacts With FZD5 and LRP5 to Regulate Proliferation and Self-Renewal of Endometrial Mesenchymal Stem-Like Cells. Front Cell Dev Biol 10:837827.

46. Owusu-Akyaw A, Krishnamoorthy K, Goldsmith LT, & Morelli SS (2019) The role of mesenchymal-epithelial transition in endometrial function. Hum Reprod Update 25(1):114–133.

47. Casazza A, et al. (2013) Impeding macrophage entry into hypoxic tumor areas by Sema3A/Nrp1 signaling blockade inhibits angiogenesis and restores antitumor immunity. Cancer Cell 24(6):695–709.

48. Wang X, et al. (2014) Functional role of arginine during the peri-implantation period of pregnancy. I. Consequences of loss of function of arginine transporter SLC7A1 mRNA in ovine conceptus trophectoderm. FASEB J 28(7):2852–2863.

49. Robertson SA, Chin PY, Femia JG, & Brown HM (2018) Embryotoxic cytokines-Potential roles in embryo loss and fetal programming. J Reprod Immunol 125:80–88.

50. Sancakli Usta C, Turan G, Bulbul CB, Usta A, & Adali E (2020) Differential expression of Oct-4, CD44, and E-cadherin in eutopic and ectopic endometrium in ovarian endometriomas and their correlations with clinicopathological variables. Reprod Biol Endocrinol 18(1):116.

51. Le Friec G, et al. (2012) The CD46-Jagged1 interaction is critical for human TH1 immunity. Nat Immunol 13(12):1213–1221.

52. Bielfeld AP, et al. (2019) A Proteome Approach Reveals Differences between Fertile Women and Patients with Repeated Implantation Failure on Endometrial Level(-)Does hCG Render the Endometrium of RIF Patients? Int J Mol Sci 20(2).

53. Feng Zhou & Roy S (2015) SnapShot: Motile Cilia. Cell 162(1):224–224.

54. Bo Li Y-PY, Yu-Ying He, Chen Liang, Meng-Yuan Li, Ying Wang, Zeng-Ming Yang (2023) IHH, SHH, and primary cilia mediate epithelial-stromalcross-talk during decidualization in mice. Sci Signal 16(774):eadd0645.

55. Li B, et al. (2022) Primary Cilia Restrain PI3K-AKT Signaling to Orchestrate Human Decidualization. International Journal of Molecular Sciences 23(24):15573.

56. Devesa-Peiro A, Sebastian-Leon P, Parraga-Leo A, Pellicer A, & Diaz-Gimeno P (2022) Breaking the ageing paradigm in endometrium: endometrial gene expression related to cilia and ageing hallmarks in women over 35 years. Hum Reprod 37(4):762–776.

57. King K, et al. (2021) Dynamics of lipid droplets in the endometrium and fatty acids and oxylipins in the uterine lumen, blood, and milk of lactating cows during diestrus. J Dairy Sci 104(3):3676–3692.

58. Vilella F, Ramirez LB, & Simon C (2013) Lipidomics as an emerging tool to predict endometrial receptivity. Fertil Steril 99(4):1100–1106.

59. Braga D, et al. (2019) Lipidomic profile as a noninvasive tool to predict endometrial receptivity. Mol Reprod Dev 86(2):145–155.

