## supporting information for "Human receptive endometrial assembloid for deciphering the implantation window"

**This PDF file includes:**

- Supplementary text
- Figures S1 to S4
- Legends for Video S1~ Video S5
- Tables S1 to S8
- Legends for Datasets S1 to S11
- SI References

**Other supporting materials for this manuscript include the following:**

- Video S1 to S5
- Datasets S1 to S11

### Supporting Materials and Methods

**Establishment of endometrial assembloids**

Endometrium was aseptically collected into pre-cooled collection medium (DMEM/F12 （Gibco，11039021）+10%FBS（Sigma，F0926）+1% antibiotic-antimycotic (Anti-Anti, Gibco, 15240-062)) stored temporarily at 4℃. After washing with prechilled DPBS (Gibco, 14190136) and removing blood clots, the endometrium was minced in 1.5 ml EP tubes and transferred to digestion medium (DMEM/F12 + 1% antibiotic-antimycotic + 0.4 mg/mL collagenase V (Sigma, C-9263) +1.25 U/mL dispase II (Sigma, D4693) + 10 µg/ml DNase I (Worthington, LS002139)). Digestion occurred at 37°C for 20 min, followed by neutralization with an equal volume of DMEM/F12 (10% FBS, 1% Anti-Anti). The digested suspension was vortexed, allowed to stand for 1 min, and filtered through a 40 µm cell strainer (Corning, 352340).

The cell strainer was inverted on the culture dish and rinsed with DPBS to collect the targeted cells. The resulting cell suspension was centrifuged at 400 × g for 5 min. The supernatant was removed, 1 ml DMEM/F12 medium was added to resuspend the cell pellet, and the pellet was centrifuged again at 400 × g for 5 min. After discarding the supernatant, the tube was chilled for 2-3 minutes, and cells were resuspended in DMEM/F12. Then, the appropriate amount of Matrigel (Corning, 536231) (volume ratio of cell suspension to Matrigel = 1:3) was added to the centrifuge tube and slowly pipetted to mix. Matrigel-cell compound was added slowly to the 24-well plate (1 drop per well, 40 µl/drop) and incubated at 37°C for 30 minutes. Expansion medium (ExM) (Table S1) (500 µL) was overlaid per well and changed every other day.

**Passage of endometrial assembloid**

Assembloids were washed twice with DPBS after medium aspiration. Digestion solution was added, mixed with cells, and incubated at 37°C for 20 min. An equal volume of neutralizing medium was added, and the cell suspension was centrifuged at 400 × g for 5 min in a 15 ml tube. After discarding the supernatant, the cell sediments were washed with DMEM/F12 medium and centrifuged at 400 × g for 5 min to acquire the sunken cells. The cell mass was resuspended in DMEM/F12, mixed with Matrigel (volume ratio of cell suspension to Matrigel = 1:3), and seeded into plates. ExM was added as previously described. The P1~P3 generation endometrial assembloids were used for downstream experiment.

**Cryopreservation of endometrial assembloids**

Assembloids were cryopreserved during each passage. The assembloid culture was rinsed with DPBS, and then the assembloid was harvested with cell recovery solution (Corning 354253) after incubating on ice for less than 30 min. Following centrifugation at 400 × g for 5 min, the assembloid was resuspended in serum-free, animal protein-free cell freezing medium (NCM Biotech, C40100), stored at -80°C overnight, and transferred to liquid nitrogen for long-term storage.

To recover the assembloids, they were removed from liquid nitrogen, thawed at 37°C, and resuspended in prewarmed DMEM/F12. The cell suspension was transferred to 15 ml tubes, diluted with additional medium, and centrifuged at 400 × g for 5 min. Cell precipitates were washed with DMEM/F12, and cell-Matrigel suspensions were prepared and seeded into plates as previously described. Y-27632 (10 μM, Millipore scm075) was added to the assembloid medium for the first three cellular fluid exchange and conventional medium thereafter.

**Long-term live-cell imaging and analysis**

Endometrial assembloids from each group were placed in the Etaluma LS720 Microscope (Bio-Reach Co., Ltd.) for continuous live-cell imaging. The growth dynamics of the assembloids were monitored, and changes in assembloid counts, area, and average intensity were recorded over time to assess their developmental progression. The average intensity reflects the growth status of the assembloids by measuring their gray value. When assembloids undergo apoptosis, they typically condense into increasingly dense solid spheres, resulting in an elevated gray value (average intensity).

**Frozen section, periodic acid-schiff staining and immunofluorescence analysis**

Assembloids were fixed in 4% PFA for 30 min, washed with PBS, dehydrated in 20% sucrose overnight at 4°C, and embedded in OCT. The embedded assembloid was sectioned using a Thermo frozen microtome at a thickness of 10 μm. The sections were applied to periodic acid-schiff staining (PAS) (Solarbio G1280).

As for immunofluorescence staining, sections were brought to room temperature, fixed with 4% PFA for 5 min, and permeabilized with 0.3% Triton X-100 in PBS for 20 min. Antigen retrieval was conducted using sodium citrate at 95°C for 20 min. After lowered to room temperature, the sections were blocked in QuickBlock™ Blocking Buffer for Immunol Staining (Beyotime Biotechnology, P0260) for 20 min. Sections were incubated with primary antibodies (Table S3) at 4°C overnight and then washed 3 times with PBS containing 0.1% Triton X-100. Secondary antibodies (Table S4) were incubated at room temperature for 2 hours, followed by three washes with 0.1% Triton X-100 in PBS. The slices were incubated with DAPI (Beyotime Biotechnology C1002) for 15 min and mounted with an anti-fluorescence quencher. The pictures were collected by confocal laser scanning microscope (Andor Dragonfly 200) and processed with Imaris x64 9.0.1.

We quantified the fluorescence images using ImageJ. Firstly, preprocess them by adjusting brightness and contrast, and removing background noise with the "Subtract Background" feature. Secondly, set the threshold to highlight the cells, then select the regions of interest (ROI) using selection tools. Thirdly, as for counting the cells, navigate to Analyze > Analyze Particles. As for measuring the influence intensity and area, set the "Measurement" options as mean gray value. Adjust parameters as needed, and view results in the "Results" window. Save the data for further analysis and ensure consistency throughout your measurements for reliable results.

DAPI was used as an internal reference for normalization, where both DAPI and target fluorescence channel intensities were quantified simultaneously. The normalized target signal intensity (target/DAPI ratio) was then compared across experimental groups. A minimum of 15 images were analyzed for each parameter per group.

**Assembloid clearing and 3D imaging**

The assembloids were fixed and cleared following Hans Clevers' Nature Protocols (1). Primary and secondary antibodies are listed in Tables S3 and S4, respectively. The images were captured by a light-sheet microscope (Carl Zeiss, Lightsheet 7) and analyzed via ZEN Blue software.

For 3D fluorescence quantification, ZEN software (Carl Zeiss) was exclusively used, with 11 images analyzed per experimental group.

**Bulk transcriptome sequencing of endometrial assembloids**

One assembloid was removed from the droplet, washed several times with DPBS and placed in lysis buffer using a capillary glass pipette under a dissecting microscope. The synthesis and amplification of full-length cDNAs were performed following the Smart-seq2 protocol (2). The total RNA of the assembloid was used as the template for the first cDNA synthesis, and Oligo dT Primer was used as the reverse transcription primer. A splice sequence was added to the 3 'end of cDNA using template-switching activity of Discover-sc Reverse Transcriptase. The RT reaction was normally performed at 42 °C for 90 min and 70°C for 15min. After the first-strand reaction, the cDNA was amplified using 9 cycles, and purified. The concentration of each library was measured by Qubit. Check the size distribution on an Agilent high-sensitivity chip. Sequencing was performed on an Illumina X-ten platform (Gene Denovo, Guangzhou, China).

**Micro proteomics of endometrial assembloids**

Assembloids were digested, collected, and washed with DPBS. At least 1 x 105 cells were transferred to low-attachment PCR tubes (Axygen, PCR-02-L-C). Downstream 4D label-free proteomic quantification was conducted by Jingjie PTM Biolab Co., Inc. (HangZhou, China). Each sample was added lysis solution, non-contact sonicated for 3 min and heated at 95℃for 10 min. When restored to room temperature, the sample was digested in 10 ng/μL trypsin overnight at 37℃. The protein solution was reduced with 5 mM dithiothreitol for 30 min at 56 °C and alkylated with 11 mM iodoacetamide for 15 min at room temperature in darkness. Then the protein was analyzed by combination of liquid chromatography and mass spectrometry. The tryptic peptides were dissolved in solvent A (0.1% formic acid, 2% acetonitrile/in water), directly loaded onto a home-made reversed-phase analytical column (25-cm length, 75/100 μm i.d.). Peptides were separated with a gradient from 6% to 24% solvent B (0.1% formic acid in acetonitrile) over 70 min, 24% to 35% in 12 min and climbing to 80% in 4 min then holding at 80% for the last 4 min, all at a constant flow rate of 450 nL/min on a nanoElute UHPLC system (Bruker Daltonics). The peptides were subjected to capillary source followed by the timsTOF Pro (Bruker Daltonics) mass spectrometry. The electrospray voltage applied was 1.75 kV. Precursors and fragments were analyzed at the TOF detector, with a MS/MS scan range from 100 to 1700 m/z. The timsTOF Pro was operated in parallel accumulation serial fragmentation (PASEF) mode. Precursors with charge states 0 to 5 were selected for fragmentation, and 10 PASEF-MS/MS scans were acquired per cycle. The dynamic exclusion was set to 30s.

**Endometrial receptivity analysis test of endometrial assembloids**

Endometrial assembloids were obtained from Matrigel via cell recovery solution and placed into RNA-later buffer (AM7020; Thermo Fisher Scientific, Waltham, MA, USA). Total RNA was extracted using RNeasy Micro Kit (Qiagen 74004). Quantification and quality were assessed by a Qubit High Sensitivity RNA Kit (Thermo Fisher Scientific Q32855) and Agilent 2100 Bioanalyzer. Assembloids with RNA integrity number (RIN) >7 were used for downstream experiments. RNA reverse transcription, cDNA library construction and endometrial receptivity determination were performed by Yikon Genomics (Jiangsu, China) (3). The MALBAC® Platinum Single Cell RNA Amplification Kit (KT110700796; Yikon Genomics, Suzhou, Jiangsu, China) was used for RNA reverse transcription. cDNA lengths of 1,000 - 10,000 bp were in accordance with the quality control requirements. Library construction was performed using the Gene Sequencing and Library Preparation Kit (XY045; Yikon Genomics, Suzhou, Jiangsu, China). Single-end sequencing was performed on a HiSeq 2500 platform (Illumina, San Diego, CA, USA). The read length was set to 140 bp. The capacity of the raw data was approximately 5M reads. The DEGs of every sample were put into the predictive model for predict the endometrial receptivity. ERA was employed specifically as a confirmatory test for the WOI assembloids, rather than as a comparative measure across all groups.

**Single-cell transcriptome sequencing and analysis of endometrial assembloids**

The CTRL, SEC and WOI endometrial assembloids from one patient were digested with digestion medium similar to digestion during passaging, but the duration of digestion was extended to 30 min. The cell precipitate obtained by centrifugation continued to be digested with 0.25% trypsin (HyClone, SH30042.01) into single cell suspensions. An equal volume of neutralizing medium was added to halt the digestion and centrifuged to acquire the cells. Cells were resuspended in PBS containing 0.1% BSA, and the cellular suspension was passed through a 40 μm cell strainer. The filtered cells were collected and subjected to cell counting and viability assays.

Cellular suspensions were loaded on a 10X Genomics GemCode Single-cell instrument that generates single-cell Gel Bead-In-EMlusion (GEMs). The resulting GEMs were processed by Gene Denovo Biotechnology Co. (Guangzhou, China). Libraries were generated and sequenced from the cDNAs with Chromium Next GEM Single Cell 3' Reagent Kits v3.1. The cellular gene matrices of each sample, produced via unique molecular identifier (UMI) counting and cell barcodes, were individually imported into Seurat version 3.1.1 for downstream analysis. Cells with an unusually high number of UMIs (≥8000) or mitochondrial gene percentage (≥10%) were filtered out. We also excluded cells with fewer than 500 or more than 4000 genes detected. Ultimately, we acquired 8608, 8932 and 9243 cells in the CTRL, SEC and WOI groups, respectively, with more than 40000 reads and approximately 2000 genes per cell after quality control (Fig. S2A). The single-cell trajectory was analyzed using a matrix of cells and gene expression by Monocle (Version 2.10.1) (4). We used CellphoneDB to analyze the expression abundance of ligand-receptor interactions between two cell types on the basis of the expression of a receptor by one cell type and a ligand by another cell type (5). Human transcription factor database (TFDB) was applied for annotating transcription factors to explore downstream gene expression and upstream epigenetic regulation in cells.

**Single-cell transcriptome analysis combined with published dataset**

The data quality of four raw datasets (CTRL, SEC and WOI assembloids, and mid-secretory endometrium) were assessed using Cellranger, followed by a comparison with the reference genome of Homo sapiens Ensembl_release103 to derive the expression matrix. Next, the expression matrices of samples were imported into Seurat. Cells were filtered using various metrics to ensure the retention of high-quality cells. This included the removal of doublets using DoubletFinder and the elimination of aberrant cells based on gene expression, UMI count, and mitochondrial gene ratio. Subsequently, we applied Harmony for batch effect correction and data integration. The dataset underwent dimensionality reduction through PCA. The soft k-means clustering algorithm was employed to address batch effects and clustering, utilizing a clustering parameter resolution of 0.5. Finally, the clustering results were visualized using tSNE or UMAP based on the cell subpopulation classification.

**Flow Cytometric Analysis**

Flow cytometry was performed to investigate the distribution of T cells and macrophages in the endometrial assembloid using post-digestive single-cell suspensions. T cells and macrophages were then stained using different staining panels (Table S5).

For staining of T cells, $7 \times 10^4$ single cells were washed by ice-cold staining buffer (PBS containing 2% fetal bovine serum), and stained for surface markers with antibodies including PerCP-Cy 5.5-conjugated anti-human CD45 antibody, V450-conjugated anti-human CD3 antibody, BV510-conjugated anti-human CD4 antibody and APC/Cyanine7-conjugated anti-human CD8 antibody (Table S5) for 30 min at 4°C in the dark, following incubating with Human TruStain FcX™ (Fc Receptor Blocking Solution) (Table S5) for 10 min at room temperature. Cells were then stained with Zombie Green™ Fixable Viability Kit (Table S5) in the dark for 20 min at room temperature.

For staining of macrophages, $7 \times 10^4$ single cells were dealt with as mentioned above except for staining for surface markers with PerCP-Cy 5.5-conjugated anti-human CD45 antibody, Brilliant Violet 785-conjugated anti-human CD68 Antibody and Brilliant Violet 650-conjugated anti-human CD11b Antibody (Table S5) and staining with Zombie NIR™ Fixable Viability Kit.

After staining, cells were fixed by 4% paraformaldehyde (PFA), washed once and then resuspended in 300 μl PBS followed by flow cytometric analysis on a BD LSR Fortessa instrument (BD Bioscience). Data were all analyzed by FlowJoV10 (BD Bioscience) in this study.

**Transmission electron microscopy**

Samples were rinsed quickly with PB buffer, immediately placed into 3% glutaraldehyde fixation solution (pH 7.4) and trimmed to 1mm x 1mm x 3mm. The following experiments were carried out at Weiya Electron Microscopy Laboratory (Jinan, Shandong, China). Rinse sequentially according to the standard TEM sample preparation methods. Then the samples were postfixed in 1% osmium tetroxide, dehydrated in series acetone, infiltrated, and embedded with Epon812. After semi-thin sectioning, use an ultramicrotome (LKB-V, LKB Company, Sweden) for 70~100nm ultrathin sectioning. The sections were stained with lead citrate and uranyl acetate and observed using a transmission electron microscope (JEM-1200EX; JEOL Ltd., Tokyo, Japan). The images were recorded using a CCD camera (MORADA-G2, Olympus Corporation, Japan).

Pinopodes are large, bulbous protrusions with a smooth apical membrane. Under transmission electron microscopy (TEM), it can be observed that the pinopodes contain various small particles, which are typically extracellular fluid and dissolved substances.

Microvilli are elongated, finger-like projections that typically exhibit a uniform and orderly arrangement, forming a "brush border" structure. Under transmission electron microscopy, dense components of the cytoskeleton, such as microfilaments and microtubules, can be seen at the base of the microvilli.

The cilium is composed of microtubules. The cross-section shows that the periphery of the cilium is surrounded by nine pairs of microtubules arranged in a ring. The longitudinal section shows that the cilium has a long cylindrical structure, with the two central microtubules being quite prominent and located at the center of the cilium.

For TEM-observed pinopodes, glycogen particles, microvilli, and cilia, manual region-of-interest (ROI) selection was performed using ImageJ software for quantitative analysis of counts, area, and length. Twenty randomly selected images per experimental group were analyzed for each morphological parameter.

**Real-Time Quantitative PCR**

Total RNA was extracted and its concentration was determined as above bulk transcriptome sequencing methods. Genomic DNA contamination was eliminated using the gDNA wiper Mix (Vazyme, R323). Total RNA (500ng) from each sample was transcribed in a total reaction volume of 20 μL using HiScript III qRT SuperMix (Vazyme, R323). Real-time PCR was performed using SYBR Green Premix Pro Taq HS qPCR Kit (Accurate Biotechnology (human) Co.,Ltd AG11701) and PCR primers (BioSune) (Table S6) in a 10 μL reaction volume. GAPDH was used as reference gene.

**Proximity ligation assay (PLA)**

The frozen sections were pretreated for fixation, permeabilized and antigen retrieval as above immunofluorescence staining. The following steps were carried out according to the Duolink® PLA Fluorescence Protocol (Sigma DUO92008, DUO82049).

The sample was blocked with Blocking Solution and incubated at 37 °C for 1 hour. Discard Blocking Solution and incubate with primary antibody (Table S7) at 4℃ overnight. On the second day, remove the primary antibody, clean with Wash Buffer A, and incubate with PLUS and MINUS PLA probe (PLUS: MINUS PLA probe = 1:5) at 37℃ for 1 hour. The sample was then incubated with Ligation Solution at 37 °C for 30min, followed by amplified with Amplification Solution at 37 °C for 100min. After washed with Wash Buffer B, the slice was sealed by Duolink® PLA Mounting Medium with DAPI. The images were captured under the confocal laser scanning microscope (Andor Dragonfly 200) and processed with Imaris x64 9.0.1.

**ATP detection**

Assembloids were digested (as described above) to obtain cell pellet. The following steps were carried out according to the ATP Test Kit (Beyotime S0026). The lysate was added to lyse the cell pellet (200μl/2*10^5 cells), followed by centrifugation at 4℃ 12000g for 5 min. The resulting supernatant was used for subsequent detection.

Preparation of standard curve: ATP standard solution of 0.01, 0.03, 0.1, 0.3, 1, 3 and 10μM can be detected to acquire the standard curve.

Preparation of ATP test working solution: Take an appropriate amount of ATP test reagent and dilute it with ATP test reagent diluent at the ratio of 1:9.

Determination of ATP concentration: Add 100μl of ATP test solution to the test hole at room temperature for 3-5 minutes to reduce the background. Add 20μl of sample or standard to the test hole and quickly mix them, which was followed by detecting the relative light units (RLU) value with the luminometer. The concentration of ATP in the sample was calculated according to the standard curve.

**Human IL-8 detection**

The medium of each-well assembloids was collected, and centrifuged at 1000 g for 10 minutes. The supernatant was put into conducted follow-up experiments according to the instructions of Human IL-8/CXCL8 ELISA Kit (ABclonal RK00011).

Wash the microwell, add 100μL standard/sample diluent (R1) into the blank well, then add 100μL standard or sample with different concentrations into the other holes. The microwell plate was incubated at 37℃ for 2 hours. Discard the liquid in the well and wash the microwell. Biotinized antibody working solution (100μL/ well) was added to each well and incubated at 37°C for 1 hour. Discard the liquid in the well and wash the microwell. Streptavidin-HRP working solution (100μL/ well) was added to each well and incubated at 37°C for 30 min. Discard the liquid in the well and wash the microwell. Add TMB substrate (100μL/ well) to the microwell, which was incubated at 37 °C for 15-20 minutes in the dark. Add the termination solution (50μL/ well), then immediately put the plate into the microplate reader and determine the OD value of 450nm within 5 minutes. The correction wavelength was set to 570 nm. Subtract the 570 nm reading from 450 nm reading to correct and remove the OD value of the non-chromogenic substance, resulting in a more accurate detection result. The concentration of IL8 for each sample is calculated according to the standard curve.

**Co-culture of blastoids and endometrial assembloids**

Blastoids were generated following the method reported in the 2021 Nature (6). The endometrial assembloids culture medium was aspirated, and pre-equilibrated modified In Vitro Culture Medium 1 (mIVC1) was added (see TableS8). On a 37°C warming plate, Day 6 blastoids were placed into endometrial assembloids using a glass needle, and immediately transferred to an incubator maintained at 37°C, 6% $CO_2$, and saturated humidity. Subsequently, half-medium changes were performed daily. At Day 8 of culture, the medium was switched to mIVC2 (see Table S8), and culture was continued until Day 9. Blastoids that remained firmly attached to endometrial assembloids during medium changes and fluorescence staining were considered to have undergone interaction with assembloids. Interaction rate = (Number of interacting embryoid-like structures / Total number of embryoid-like structures) × 100%.

All biological replicates (fourteen individuals) of endometrial assembloids show similar expression of receptivity markers and comparable capacity to support blastoid attachment.
