## Supplementary material for "Human receptive endometrial assembloid for deciphering the implantation window": Video_Table_Dateset

**Video S1** Representative video of the endometrial glands gradually developing into a vesicular shape, and the surrounding stromal cells arranging in a fibrous pattern in the endometrial assembloids, as imaged at 100x by time-lapse microscopy of KEYENCE BZ-X800E over 72 h.

**Video S2** Representative video of the stromal cells growing in fibrous pattern and forming an extensive network in the endometrial assembloids, as imaged at 200x by time-lapse microscopy of KEYENCE BZ-X800E over 48 h.

**Video S3~S4** Representative video showing that the endometrial assembloids grew and differentiated, the endometrial glands enlarged, and surrounding stromal cells formed an extensive network during the hormone treatment (WOI) and control environment (CTRL), as imaged at 100x by time-lapse microscopy of KEYENCE BZ-X800E over 8 days.

**Video S5** Representative video showing that three-dimensional imaging of the chemically cleared CTRL and WOI assembloids, as imaged by light-sheet microscope to detect the expression of FOXO1 and FOXA2.

**Table S1. Composition of expansion medium (ExM) of endometrial assembloid**

| Reagent | Manufacturer | Catalog no. | Concentrations |
|---|---|---|---|
| DMEM/F12 | Gibco | 11039-021 | |
| Antibiotic-Antimycotic (100X) | Gibco | 15240062 | 1% |
| ITS | Gibco | 41400-045 | 1% |
| L-Glutamine | Gibco | 25030-081 | 2 mM |
| Nicotinamide | Sigma | N3376 | 1 mM |
| B27 | Gibco | 17504-044 | 2% |
| N2 | Gibco | 17502-048 | 1% |
| Noggin | Proteintech | HZ-1118 | 100 ng/ml |
| EGF | Peprotech | AF-100-15 | 50 ng/ml |
| FGF2 | Origene | TP750002 | 100 ng/ml |
| WNT-3A | Proteintech | HZ-1296 | 200 ng/ml |
| R-Spondin-1 | Peprotech | 120-38 | 200 ng/ml |
| A83-01 | MCE | HY-10432 | 0.5 uM |
| N-acetyl-L-cysteine | Sigma | A7250 | 1.25 mM |
| p38 inhibitor SB202190 | Sigma | SB202190 | 10 uM |

**Table S2. Composition of hormone regimen of endometrial assembloid**

| Reagent | Manufacturer | Catalog no. | Concentrations |
|---|---|---|---|
| Estradiol | Sigma | E2758 | 10nM |
| Medroxyprogesterone Acetate | Selleck | S2567 | 1μM |
| N6,2′-O-dibutyryladenosine 3′,5′-cyclic monophosphate sodium salt (cAMP) | Sigma | D0627 | 1μM |
| Human Chorionic Gonadotropin (HCG) | Livzon Pharmaceutical Group Inc | 2000U | 1μg/ml |
| Human Placental Lactogen (HPL) | R&D Systems | 5757-PL | 20ng/ml |
| Prolactin | Peprotech | 100-07 | 20ng/ml |

**Table S3. Primary antibodies used in immunofluorescence**

| Antibody | Manufacturer | Catalog no. | Dilution ratio |
|---|---|---|---|
| E-Cadherin (4A2) Mouse mAb | Cell Signalling | 14472 | 1:200 |
| Vimentin (D21H3) XP® Rabbit | Cell | 5741S | 1:200 |

| | | | |
|---|---|---|---|
| mAb | Signalling | | |
| Ki-67 (D3B5) Rabbit mAb | Cell Signalling | 9129 | 1:400 |
| Rabbit anti-Cleaved Caspase-3 | Cell Signalling | 9661S | 1:400 |
| Progesterone Receptor A/B (D8Q2J) XP® Rabbit mAb | Cell Signalling | 8757S | 1:800 |
| FoxO1 (D7C1H) Mouse mAb | Cell Signalling | 14952S | 1:100 |
| Rabbit anti-FOXA2 | Abcam | ab108422 | 1:500 |
| Anti-PDGF B 抗体 | Abcam | ab23914 | 1:160 |
| Anti-Integrin alpha 5 | Abcam | ab150361 | 1:222 |
| Anti-CD44 | Abcam | ab254530 | 1:500 |
| Mouse anti-Estrogen Receptor alpha | Santa Cruz | sc-8002 | 1:50 |
| neuropilin-2 antibody | Santa Cruz | sc-13117 | 1:50 |
| TPPP antibody | Santa Cruz | sc-515819 | 1:50 |
| Goat anti-Human DPPIV/CD26 | R&D Systems | AF1180 | 1:100 |
| HIF-1 alpha antibody | NOVUS | NB 100-105 | 1:50 |
| Acetyl-α-tubulin | Sigma | T7451 | 1:20000 |
| Anti-ODF2 antibody | Sigma | HPA 048841 | 1:100 |
| SLC25A1 Polyclonal antibody | Proteintech | 15235-1-AP | 1:500 |
| CD45 Monoclonal Antibody (30-F11) | eBioscience | 14-0451-81 | 1:100 |
| Anti-IGFBP1 | Abcam | ab228741 | 1:400 |
| MAO-A | Santa Cruz | sc-271123 | 1:50 |

**Table S4. Second antibodies used in immunofluorescence**

| Antibody | Manufacturer | Catalog no. | Dilution ratio |
|---|---|---|---|
| Donkey anti-Mouse IgG (H+L) Highly Cross-Adsorbed Secondary Antibody Alexa Fluor™ Plus 488 | Thermo | A32766 | 1:1000 |
| Donkey anti-Mouse IgG (H+L) Highly Cross- | Thermo | A32773 | 1:10000 |

| | | | |
|---|---|---|---|
| Adsorbed Secondary Antibody Alexa Fluor™ Plus 555 | | | |
| Donkey anti-Rabbit IgG (H+L) Highly Cross-Adsorbed Secondary Antibody Alexa Fluor™ Plus 555 | Thermo | A32794 | 1:1000 |
| Donkey anti-Rabbit IgG (H+L) Highly Cross-Adsorbed Secondary Antibody Alexa Fluor™ Plus 488 | Thermo | A32790 | 1:1000 |
| Donkey anti-Rabbit IgG (H+L) Highly Cross-Adsorbed Secondary Antibody Alexa Fluor™ Plus 647 | Thermo | A32795 | 1:1000 |
| Donkey anti-Goat IgG (H+L) Highly Cross-Adsorbed Secondary Antibody Alexa Fluor™ Plus 647 | Thermo | A32849 | 1:1000 |
| Goat anti-Rat IgG (H+L) Cross-Adsorbed Secondary Antibody, Alexa Fluor™ 488 | Thermo | A11006 | 1:500 |

**Table S5. Antibodies and related reagents used in flow cytometry**

| Antibody | Manufacturer | Catalog no. | Dilution ratio |
|---|---|---|---|
| BD Pharmingen™ PerCP-Cy™5.5 Mouse Anti-Human CD45 | BD Bioscience | 564105 | 1:200 |
| BD Horizon™ V450 Mouse Anti-Human CD3 | BD Bioscience | 560365 | 1:200 |
| BD Horizon™ BV510 Mouse Anti-Human CD4 | BD Bioscience | 562970 | 1:200 |
| APC/Cyanine7 anti-human CD8 Antibody | Biolegend | 344714 | 1:200 |
| CD127 Monoclonal Antibody (eBioRDR5), Alexa Fluor™ 700 | invitrogen | 56-1278-42 | 1:200 |
| Brilliant Violet 785™ anti-human CD68 Antibody | Biolegend | 333825 | 1:200 |
| Brilliant Violet 650™ anti-human CD11b Antibody | Biolegend | 301335 | 1:200 |
| Human TruStain FcX™ (Fc | Biolegend | 422301 | 1:200 |

| | | | |
|---|---|---|---|
| Receptor Blocking Solution) | | | |
| Zombie Green™ Fixable Viability Kit | Biolegend | 423111 | 1:500 |
| Zombie NIR™ Fixable Viability Kit | Biolegend | 423105 | 1:500 |

**Table S6. PCR primers used in real-time quantitative PCR analysis**

| Gene | Sequence (5'to3') |
|---|---|
| PGR | F: ACCCGCCCTATCTCAACTACC |
| | R: AGGACACCATAATGACAGCCT |
| PAEP | F: GAGATCGTTCTGCACAGATGG |
| | R: CGTTCGCCACCGTATAGTTGAT |
| OLFM4 | F: ACCTTTCCCGTGGACAGAGT |
| | R: TGGACATATTCCCTCACTTTGGA |
| ESR1 | F: CCCACTCAACAGCGTGTCTC |
| | R: CGTCGATTATCTGAATTTGGCCT |

**Table S7. Antibodies used in proximity ligation assay (PLA)**

| Antibody | Manufacturer | Catalog no. | Dilution ratio |
|---|---|---|---|
| SEMA3A antibody (C-1) | Santa Cruz | sc-74555 | 1:50 |
| Recombinant Anti-Neuropilin 1 antibody [EPR3113] | Abcam | ab81321 | 1:250 |
| Recombinant Anti-ROR2 antibody [EPR19980] | Abcam | ab218105 | 1:500 |
| Wnt-5a antibody (A-5) | Santa Cruz | sc-365370 | 1:50 |
| Monoclonal Anti-CD74 antibody produced in mouse | Sigma | SAB5201932 | 1:100 |
| Recombinant Anti-alpha COP I/COPA Antibody [EPR14273(B)] | Abcam | ab181224 | 1:50 |
| CD46 antibody (M177) | Santa Cruz | sc-52647 | 1:50 |
| Jagged1 (28H8) Rabbit mAb | Cell Signaling | 2620T | 1:1000 |

**Table S8. Composition of modified In Vitro Culture Medium (mIVC1 and mIVC2) for co-culture of blastoids and endometrial assembloids**

| Reagent | Manufacturer | Catalog no. | Concentrations |
|---|---|---|---|
| mIVC1 | | | |
| advanced DMEM/F12 | Gibco | 12634-010 | - |
| defined fetal bovine serum | Biosera | bs-0003 | 20% |
| L-glutamine | Gibco | 25030 | 2 mM |
| ITS-X | Gibco | 51500-056 | 1x |
| β-estradiol | Sigma | E8875 | 8 nM |
| progesterone | Sigma | P0130 | 200 ng/ml |
| N-acetyl-L-cysteine | Sigma | A7250 | 25 µM |
| sodium lactate | Sigma | L7900 | 0.22% |
| Sodium pyruvate | Sigma | P4562 | 1 mM |
| Y27632 | Selleck | S1049 | 10µM |
| mIVC2 | | | |
| advanced DMEM/F12 | Gibco | 12634-010 | - |
| KOSR | Gibco | A3181501 | 30% |
| L-glutamine | Gibco | 25030 | 2 mM |
| ITS-X | Gibco | 51500-056 | 1x |
| β-estradiol | Sigma | E8875 | 8 nM |
| progesterone | Sigma | P0130 | 200 ng/ml |
| N-acetyl-L-cysteine | Sigma | A7250 | 25 µM |
| sodium lactate | Sigma | L7900 | 0.22% |
| Sodium pyruvate | Sigma | P4562 | 1 mM |
| Y27632 | Selleck | S1049 | 10µM |

**Dataset S1** Clinical information of patients providing endometrial tissue.

**Dataset S2** Differentially expressed receptivity-related genes across nine hormone regimens (Corresponding to Figure 1E).

**Dataset S3** Comparison of gene expression among CTRL, SEC and WOI assembloid and early-mid secretory endometrium at single-cell level (Corresponding to Figure 2G).

**Dataset S4** Differentially expressed TFs of endometrial assembloids and early-mid secretory endometrium as for secretory epithelium and EMT-derived cells (Corresponding to Figure 2H).

**Dataset S5** Differentially expressed genes among all clusters in the assembloids at the single-cell level (Corresponding to Figure S2G~H).

**Dataset S6** Differentially expressed genes among the CTRL, SEC and WOI assembloids from Smart-seq2 transcriptome (Corresponding to Figure 3B).

**Dataset S7** Differentially expressed genes in the SEC and WOI assembloids at the single-cell level (Corresponding to Figure 3F, 5E, S2I, S4A).

**Dataset S8** Differentially expressed genes and pathways identified by Mfuzz trend analysis across CTRL, SEC, and WOI groups (Corresponding to Figure 4A).

**Dataset S9** Differentially expressed proteins among the SEC and WOI groups (Corresponding to Figure 4B).

**Dataset S10** Differentially expressed cilia-related genes across CTRL, SEC, and WOI groups (Corresponding to Figure 5A).

**Dataset S11** Ligand-receptor interactions among all clusters in the CTRL, SEC and WOI assembloids at the single-cell level (Corresponding to Figure 5G, S4E~F).

### SI References


1. J. F. Dekkers *et al.*, High-resolution 3D imaging of fixed and cleared organoids. *Nat Protoc* **14**, 1756-1771 (2019).
2. S. Picelli *et al.*, Full-length RNA-seq from single cells using Smart-seq2. *Nat Protoc* **9**, 171-181 (2014).
3. A. He *et al.*, The role of transcriptomic biomarkers of endometrial receptivity in personalized embryo transfer for patients with repeated implantation failure. *J Transl Med* **19**, 176 (2021).
4. C. Trapnell *et al.*, The dynamics and regulators of cell fate decisions are revealed by pseudotemporal ordering of single cells. *Nat Biotechnol* **32**, 381-386 (2014).
5. M. Efremova, M. Vento-Tormo, S. A. Teichmann, R. Vento-Tormo, CellPhoneDB: inferring cell-cell communication from combined expression of multi-subunit ligand-receptor complexes. *Nat Protoc* **15**, 1484-1506 (2020).
6. Yu L, et al., Blastocyst-like structures generated from human pluripotent stem

cells. Nature 591(7851):620-626 (2021).
